# Deciphering the mechanism of protein aggregation and effects of inhibitors using single-molecule mass photometry

**DOI:** 10.1101/2025.08.11.668757

**Authors:** Aaron Lyons, Simanta Sarani Paul, Jack E. MacArthur, Grace N. Hoffman, Allan Yarahmady, Sue-Ann Mok, Michael T. Woodside

## Abstract

Aggregation of misfolded proteins is a prominent feature of many diseases and hence an attractive drug target. However, the small oligomers that are critical early species in the aggregation cascade are difficult to monitor directly owing to their heterogeneity and transience, complicating efforts to define aggregation mechanisms and target oligomers therapeutically. Here, we observe changes in oligomer populations directly using single-molecule mass photometry (SMMP). Studying the pathogenic P301L mutant of tau protein linked to frontotemporal dementia, we globally fit the growth/decay kinetics for every oligomer population observed by SMMP to microscopic models of aggregation. A simple extension to the best-fit model also accounts for amyloid fibril kinetics, as monitored by Thioflavin T fluorescence, providing the first quantitative model of aggregation kinetics across all stages of the cascade based on direct observation. Crucially, we find that models fitting amyloid kinetics alone fail to capture oligomer behavior, implying that—contrary to standard practice—amyloid kinetics cannot be relied on to deduce aggregation mechanisms. Furthermore, there is no single rate-limiting nucleation step preceding rapid growth, as generally assumed, suggesting that standard models of aggregation are overly simplistic. Repeating the analysis in the presence of aggregation inhibitors allows identification of the discrete steps in the cascade affected by the inhibitors. This work presents a powerful new approach for defining protein aggregation mechanisms and the mechanism of action of inhibitors, with applications to understanding many diseases and developing novel therapeutics.

---

Aggregates of misfolded protein feature prominently in a wide range of diseases: neurodegenerative diseases (*e*.*g*. Alzheimer’s, Parkinson’s, frontotemporal dementia, amyotrophic lateral sclerosis, and prion diseases), systemic amyloidoses, type II diabetes, cardiac amyloidoses, and many others [1, 2]. Although the proteins involved in these diseases may differ, their aggregation follows a cascade with broadly similar molecular features. In general, initial misfolding or destabilization of the native structure is thought to lead to nucleation of various oligomers that proceed to grow in size, maturing ultimately into stable amyloid fibrils [3]. In some cases, secondary processes such as nucleation of oligomers from larger aggregates and fragmentation of large aggregates into smaller pieces are also proposed to play key roles [1, 3]. Given the relevance of protein aggregation to disease, it has been a major target of drug development efforts seeking inhibitors that can act as disease-modifying or preventive therapeutics [4,5]. However, these efforts have been impeded by difficulties in determining aggregation mechanisms so as to define molecular targets for inhibiting aggregation, as well as challenges in determining the mechanism of action of inhibitors.

A key challenge posed by aggregation relates to small oligomers: they are important players both as essential intermediates in the aggregation cascade and also as the molecular species thought in many cases to be most toxic to cells, particularly in neurodegenerative diseases [6]. However, they are very difficult to study, because they are not only heterogeneous in size and/or conformation but also typically present transiently as only a minor subpopulation in the aggregation reaction. Some methods used to monitor aggregation at the ensemble level are sensitive to small oligomers, such as ion mobility spectrometry-mass spectroscopy [7] and micro free-flow electrophoresis [8], but their ability to distinguish or quantify different oligomeric species is limited. In contrast, more commonly used methods are typically insensitive to minority sub-populations, leaving most of the early oligomers undetected or poorly characterized. For example, amyloid-binding dyes like Thioflavin T (ThT) are widely used to study protein aggregation and test potential inhibitors [9], inferring events in the early stages of aggregation based on the lag time preceding amyloid detection. Such measurements are often fit to kinetic models that purport to describe microscopic features of the aggregation mechanism [10, 11]. However, despite their widespread application in studies of protein aggregation and drug discovery efforts, the reliability of these models is unclear because they are not based on direct measurements of the small oligomers that play crucial roles in defining aggregation mechanisms.

Single-molecule mass photometry (SMMP) can overcome these challenges to provide a more complete description of the aggregation cascade. The mass of individual molecules in solution is quantified from the optical contrast arising from interference between back-scattered and reflected light as molecules bind non-specifically to a microscope coverslip (Fig. 1) [12], allowing the populations of each oligomer size to be quantified directly. Like other single-molecule (SM) approaches, SMMP is sensitive to sub-populations of rare or transiently occupied states that would otherwise remain hidden within ensemble averages [13], but it avoids the technical constraints (*e*.*g*. reporter labels, low throughput) that have limited the use of methods like SM imaging [14], fluorescence [15], and force spectroscopy [16]. Because the growth and decay of populations for a wide range of oligomer sizes can be observed over time, mechanistic models of aggregation can be tested directly and the microscopic rates for each step in the aggregation cascade can be deduced [17]. Previous work with SMMP has studied oligomers formed by proteins including bovine serum albumin [18], peroxiredoxins [19], and MinDE [20]. SMMP has also been used to monitor the growth of actin and α-synuclein fibrils [18], and the early stages of aggregation of wild-type tau protein [17]. However, SMMP assays have never been used to model the complete aggregation cascade, from small oligomers up to fibrils. Moreover, SMMP has never been compared directly to more standard assays like ThT fluorescence to assess which technique can more effectively describe the earliest stages of aggregation or the molecular mechanisms of aggregation inhibitors.

**Figure 1:**
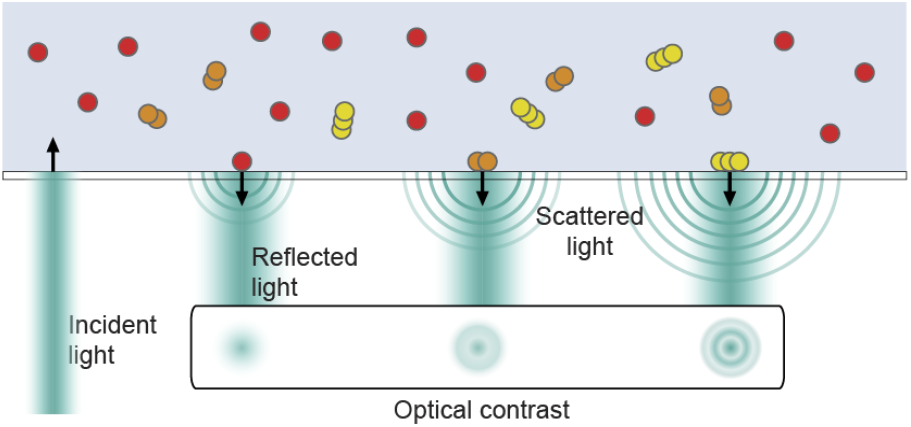
Single-molecule mass photometry of tau P301L. Schematic describing the operating principles of SMMP. Protein molecules binding to a coverslip surface scatter light in a size-dependent way. Interference with the incident light reflected from the coverslip produces optical contrast that varies with the size of the detected protein.

Here we do so for the first time, studying the aggregation of tau protein. Misfolded tau aggregates feature in multiple neurodegenerative diseases, including Alzheimer’s, frontotemporal dementia, chronic traumatic encephalopathy, and progressive supranuclear palsy [21], making tau aggregation an important diagnostic and therapeutic target. Focusing on the pathogenic P301L mutant linked to frontotemporal dementia, which unlike wild-type (WT) tau aggregates spontaneously without posttranslational modifications or inducers [22], we use SMMP to quantify the time evolution of all oligomers up to 25- mers. Identifying a minimal microscopic model consistent with the data by globally fitting all oligomer population dynamics, we find that this model also fits the kinetics of fibrils observed by ThT fluorescence. In contrast, standard kinetic analysis of ThT fluorescence alone fails to infer the oligomer population dynamics seen by SMMP, implying that assays sensitive only to fibrils do not reliably capture oligomer kinetics. Finally, we show that epigallocatechin-3-gallate (EGCG), a broad-spectrum anti-amyloidogenic compound [23], arrests aggregation by greatly reducing the growth rates of all oligomers, whereas methylene blue, a tau aggregation inhibitor previously tested clinically against Alzheimer’s disease [24], suppresses aggregation mainly by creating a bottleneck at tetramers but does not prevent the slow accumulation of larger oligomers. This work shows how SMMP can be used not only to decipher aggregation mechanisms, but also to define key steps in the aggregation cascade suitable for therapeutic intervention and to determine how inhibitors act on the cascade, highlighting the promise of SMMP for helping to develop new therapeutics for misfolding diseases.

## RESULTS

### Time-evolution of tau P301L oligomer populations observed by SMMP

We first examined the aggregation of the P301L mutant of tau in the absence of any inducers, studying the 0N4R isoform of human tau. As previously reported, spontaneous aggregation of tau P301L is quite slow, taking place over many days [22]. At 0 hr of aggregation, the mass distribution observed by SMMP (Fig. 2A) showed three peaks corresponding to monomers, dimers, and trimers, just as was seen previously for WT tau before aggregation [17]. Relative occupancies of the dimer and trimer were lower than for WT tau, implying that these initial oligomers were less stable for P301L tau than WT. Objects at higher mass (reflecting impurities or higher-order tau oligomers) were observed very infrequently, with none seen above 400 kDa (inset).

**Figure 2:**
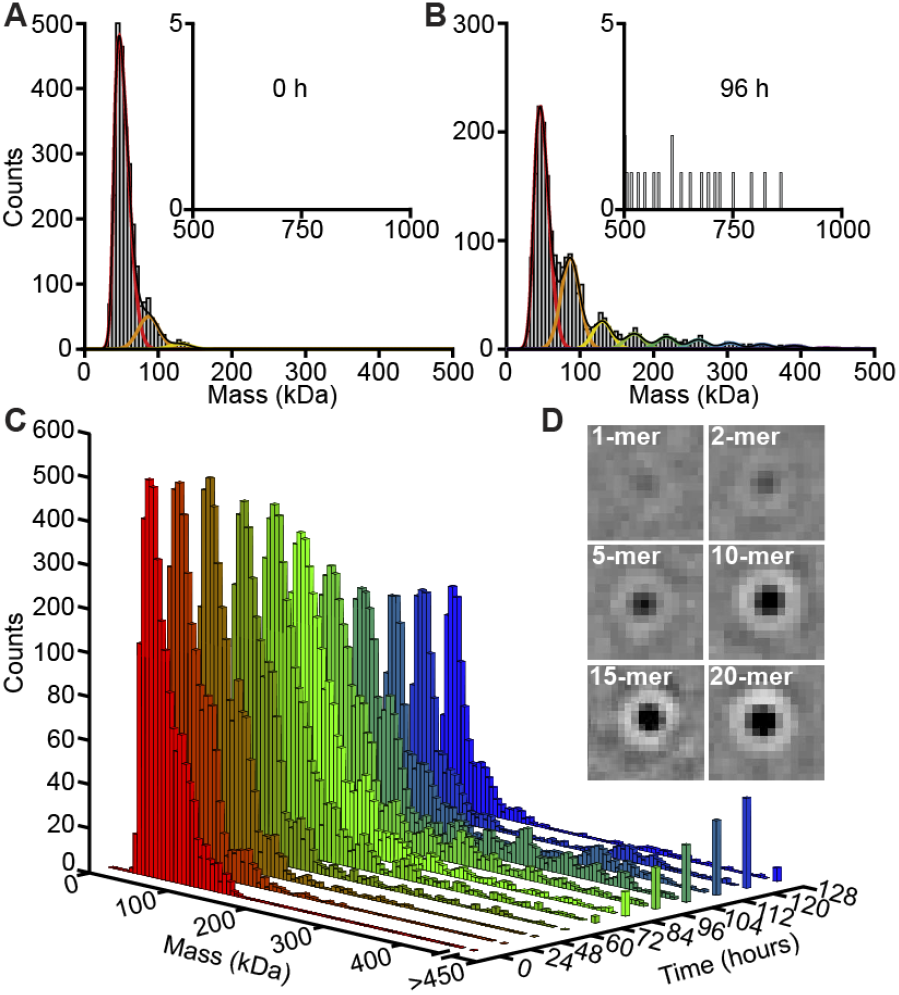
SMMP of P301L tau aggregation. (A, B) Mass distributions after (A) 0 and (B) 96 hr of aggregation. Mostly monomers are seen at 0 hr (A), with minor peaks corresponding to dimers and trimers. Monomer prevalence is lower after 96 hr (B), but now mass peaks corresponding to oligomers up to 10-mers are seen, as well as isolated events at masses up to the MDa range (inset). (C) Time evolution of oligomer populations shows larger aggregates increase in number as the aggregation cascade progresses. (D) Sample images of oligomers ranging from monomers to 20-mers. All images 1.43 × 1.43 μm, using the same contrast scale.

To quantify the time evolution of the oligomer populations during the aggregation cascade, samples were taken every ∼8 or 12 hr from a stock of tau P301L aggregating at 10 µM and diluted for measurement by SMMP. The oligomer mass distribution gradually changed from the initial result, reducing the monomer counts while gaining new peaks at masses corresponding to higher-order oligomers, with events observed up to the MDa range (Fig. 2B). As the cascade progressed, the initial decline in monomers moderated, dimer and trimer populations rose after a brief lag phase before dropping sharply, and ever-larger oligomers accumulated at ever-later times before each declining in turn (Fig. 2C). Distinct peaks could be observed consistently for oligomers up to ∼10-mers, whereas oligomers up to 25-mers could be readily identified (Fig. 2D) but at insufficient counts to produce well-defined peaks. Very large aggregates (possibly fibrillar) were also observed towards the end of the measurement, but they could not be imaged reliably to determine their mass from the light scattered. Because SMMP is typically insensitive to conformation, oligomers of the same mass but different conformations could not be distinguished.

We analyzed the time-evolution of the oligomer populations similar to previous work [17]: at each time point, we fit the areas of the peaks corresponding to monomers through 10-mers to find the total counts for each species; for larger oligomers, where clear mass peaks could not be identified, we combined the raw counts of 11-mers to 15mers into one population and the counts from 16-mers to 25-mers into a second to obtain sufficient statistics. The total counts for each oligomeric species were then plotted as a function of time, correcting for the systematic undercounting of larger oligomers owing to size-dependent diffusion (Fig. 3A).

**Figure 3:**
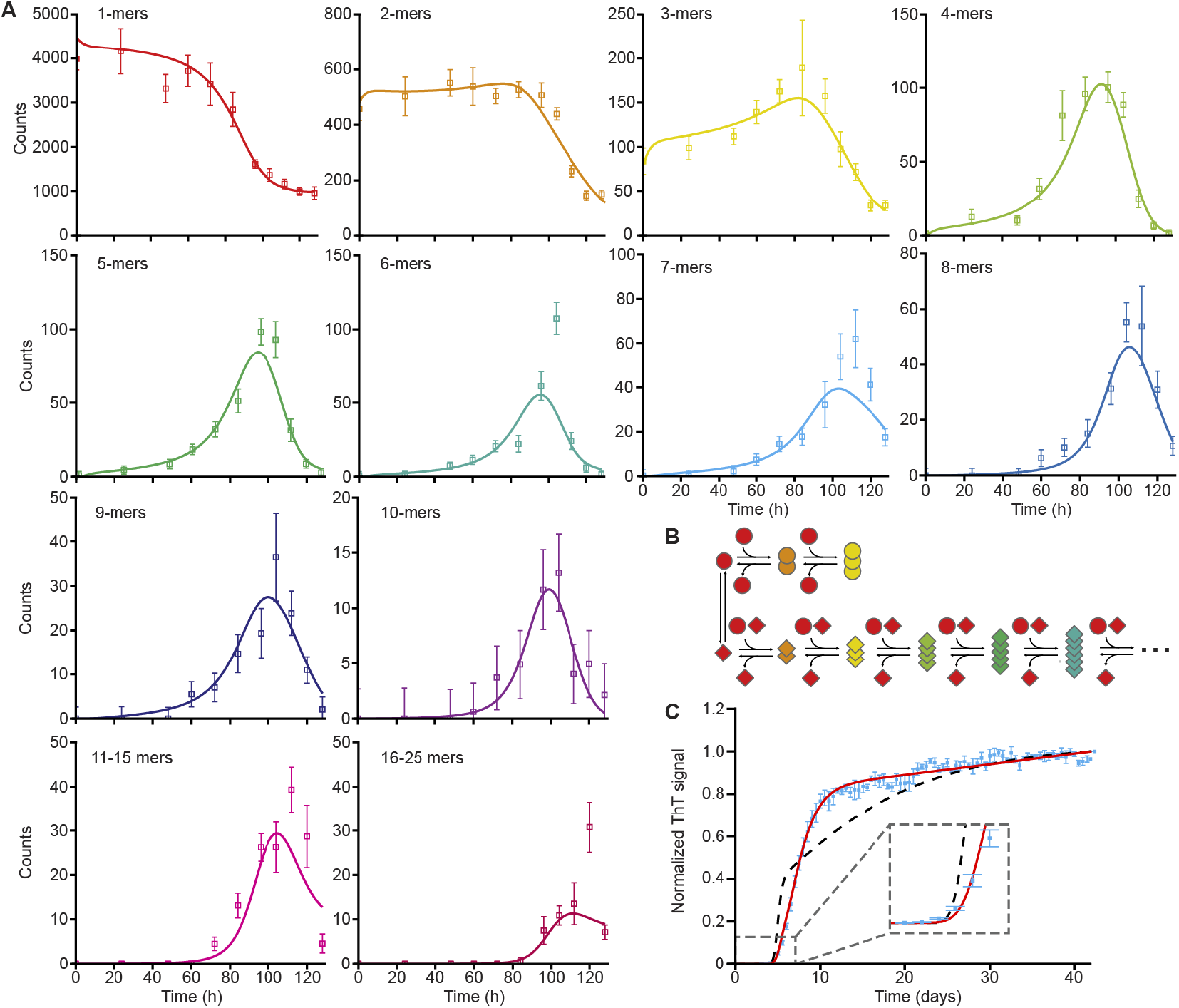
Modeling P301L tau aggregation. (A) Time evolution of population counts for monomers through 25-mers; 11–15-mers are grouped into a single population to ensure sufficient counts, as are 16–25-mers. Error bars show standard error of mean from 3 replicates. Solid lines: global fits to model in (B). (B) Minimal model of aggregation: parallel pathways for native oligomers (top, capped at trimers) and misfolded oligomers (bottom, unbounded growth); monomers add to misfolded oligomers in either native or misfolded conformers but disassociate only in misfolded conformers. (C) Kinetics of ThT fluorescence from large oligomers (cyan) are predicted reasonably well by simply projecting the fit from panel A to large oligomers (black), and are fully accounted by this model when assuming monomer addition/removal rate is different for ThT positive aggregates (red).

### Aggregation mechanism from oligomer kinetics predicts ThT fluorescence

Aggregation kinetics can be fit to microscopic kinetic models to gain insight into the mechanism of aggregation. Typically, the kinetics of large aggregates such as those assayed by ThT fluorescence are used [25], because the populations of individual oligomeric species are hard to quantify directly. However, the comprehensive information about oligomer time-evolution provided by SMMP allows highly detailed models of aggregation to be built, returning the microscopic rates for each step in the cascade [17]. It was previously found for WT tau that two parallel pathways—one involving natively folded tau and a second involving misfolded tau—were needed to account for the lag phase in the growth of dimers and trimers despite the presence of dimers and trimers prior to aggregation [17]. Because a one-pathway model was unable to account for the population dynamics observed for P301L tau (Fig. S1, Table S1), we postulated a similar two-pathway model, assuming a spontaneous equilibrium between two conformers of P301L tau (a misfolded conformer able to aggregate and a native conformer unable to form oligomers larger than trimers) and changes in oligomer size via monomer addition or subtraction (Fig. 3B). Fitting the coupled differential equations describing all oligomer population dynamics (see SI Methods) simultaneously to all oligomer populations (Fig. 3A, solid lines; Table S2), we found that this model captured almost all the essential features of the oligomer populations, although it consistently underestimated the peaks of most higher-order oligomers.

To test this model further, we projected the kinetics to larger oligomers, assuming that aggregates larger than 25mers continued to grow/shrink by monomer addition/subtraction with the same rates as for 25-mers. We then compared the model to ThT fluorescence data measured at the same tau concentration (Fig. 3C, cyan). Aggregates were assumed to become sensitive to ThT with a sigmoidal size-dependence (since small oligomers are ThT-negative whereas ThT-positive fibrils dominate among large aggregates), treating the sigmoid shape as a fitting parameter. This projection (Fig. 3C, black; Table S3) captured the timing of the growth phase well (inset) but predicted a more gradual transition to the plateau region than observed. Because ThT positivity is associated with a conformational change, which could well lead to different rates for monomer addition/removal, to provide a more realistic model we allowed these rates to differ from ThT-negative oligomers; for simplicity, we assumed all ThT-positive aggregates had the same on/off rates. This simple extension of the SMMP kinetic model matched the ThT data extremely well at all times (Fig. 3C, red; Table S4). The kinetics measured from SMMP were thus also able to account for the kinetics of both small oligomers and larger, fibrillar aggregates, with only a simple extension of the model bridging early and late-stage aggregate behavior.

We note that including extra steps in the model such as addition/removal of misfolded dimers or direct misfolding of native dimers and trimers did not improve the fits. Significantly, there was no need to include processes like fibril fragmentation or secondary nucleation (formation of oligomers stimulated by the presence of fibrils) to account for the observed fibril kinetics, even though previous work has suggested such processes are essential for explaining results from ThT fluorescence [26].

### Fibril kinetics fail to capture oligomerization

To compare the results from SMMP to those from more standard probes of aggregation mechanisms in greater detail, we applied state-of-the-art methods for analyzing the kinetics of concentration-dependent ThT fluorescence signals [10, 25]. These methods have been widely used to deduce aggregation mechanisms and effects of inhibitors [27, 28, 29]. Although previous work found that such modeling can be consistent with bulk measurements of soluble oligomers [30, 31], consistency with the detailed oligomeric cascade observed by SMMP has not yet been tested. Fitting the concentration-dependent ThT signal with Amylofit to test different aggregation mechanisms [25], we found that a simple nucleation-elongation model—analogous to the model used to fit SMMP data—fit poorly (Fig. 4A). A model featuring saturating elongation and fragmentation captured the observed concentration-dependence and long lag phases best among available models (Fig. 4B), with saturating elongation plus secondary nucleation performing only slightly worse. This need for secondary processes to describe fibril growth kinetics is consistent with previous work on a 77-aa tau fragment [32]. However, in contrast to the extended SMMP model, all Amylofit models consistently underestimated the lag time, predicting a gentler transition to the growth phase than actually observed.

**Figure 4:**
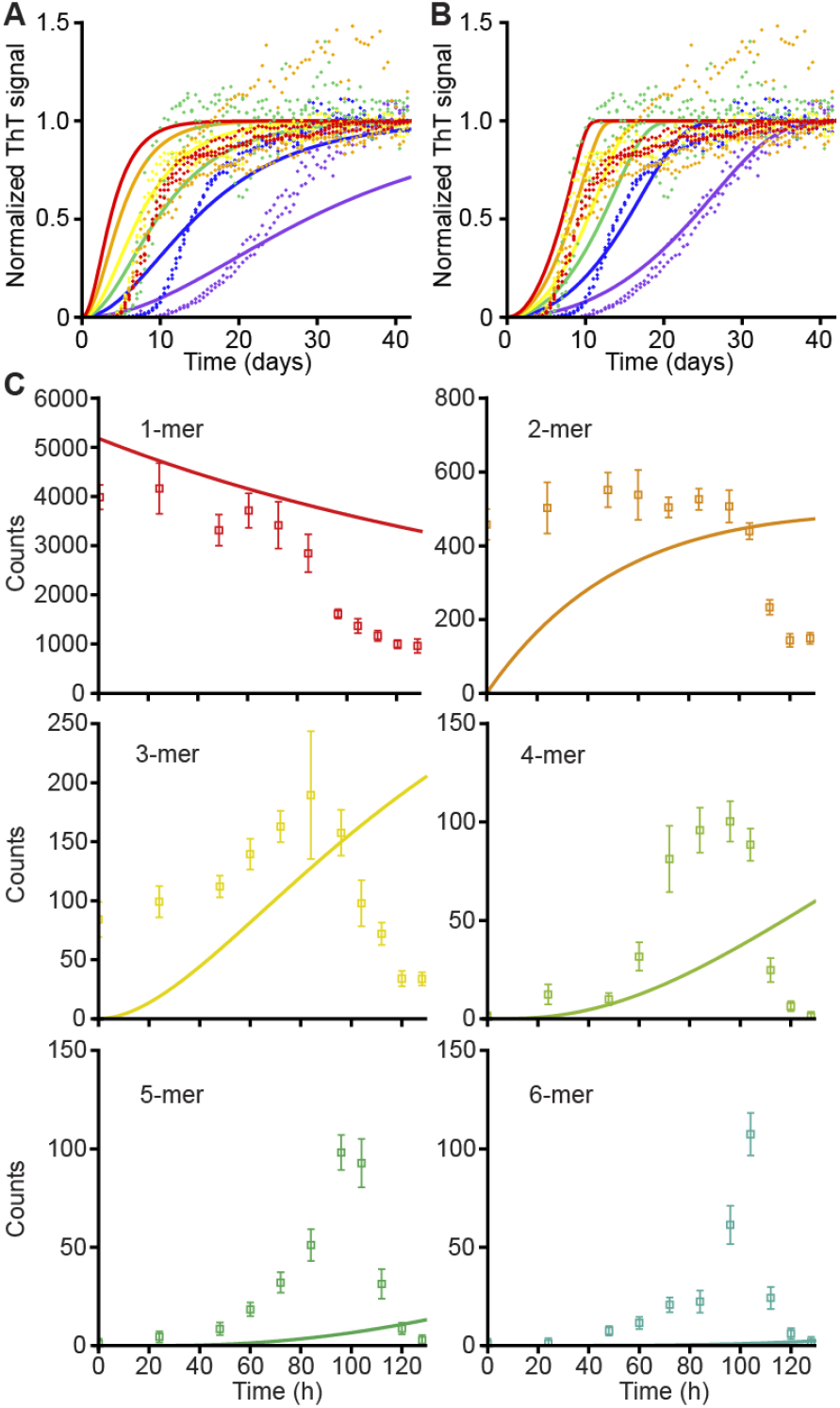
Modeling fibril kinetics fails to predict oligomer kinetics. (A, B) Concentration-dependent ThT signal fit to (A) nucleation-elongation and (B) saturating elongation with fragmentation; the latter was the best fit among all Amylofit models. (C) Oligomer dynamics predicted from best fit in (B) do not match counts observed by SMMP.

We also explored whether the best-fitting model of the ThT data predicted oligomer populations reliably. To do so, we converted the fibril model from Amylofit into an equivalent system of differential equations featuring saturating elongation and fragmentation that accounted for each oligomer population independently (SI Methods). The resulting model failed to match the oligomer populations observed by SMMP (Fig. 4C), predicting a delayed but exaggerated growth of the smallest oligomers while greatly underestimating the growth of larger oligomers (pentamers and up, Fig. S2), likely owing to the rapid fragmentation of these species. Intriguingly, repeating this process for the simple nucleation-elongation model produced a better fit (Fig. S3) despite the poor performance when modeling the ThT data. This result shows that the mechanism and kinetics inferred from global fitting of tau P301L fibrillization are inconsistent with the observed evolution of oligomer populations, implying more generally that optimizing models to fit ThT data does not necessarily yield an accurate picture of the early stages of the aggregation cascade.

### Inhibitors modulate the rates of specific steps in the aggregation cascade

Determining the mechanism of action of aggregation inhibitors is important for developing them as drug candidates. Numerous studies have explored such mechanisms via the effects of inhibitors on fibril kinetics, inferring inhibition of specific processes in the aggregation mechanisms deduced from fits [33, 34], and a handful have explored effects on ensembles of smaller oligomers [7, 35, 36]. However, the effects of inhibitors on each distinct small oligomer have never been measured directly. We therefore did so using SMMP, comparing the time evolution of oligomer populations seen for P301L tau alone with that seen when adding small-molecule inhibitors: EGCG, a catechin known to inhibit the aggregation of multiple proteins including tau, amyloid-β, and α-synuclein [23]; and methylene blue, an aggregation inhibitor that was tested as an anti-Alzheimer’s therapeutic but failed to show efficacy in late-stage clinical trials [24].

In the presence of EGCG, the three initial peaks corresponding to monomers, dimers, and trimers seen in the absence of inhibitor remained mostly unchanged over the course of six days: no clear peaks corresponding to tetramers or larger oligomers appeared, suggesting a near-complete arrest of the aggregation cascade (Fig. 5A). Quantifying the oligomer populations as before but combining the data from 4-mers through 25-mers because of low counts, we fit the resulting oligomer dynamics to the kinetic model from Fig. 3B (Fig. S4). The rates for monomer addition to all oligomer sizes were reduced by several orders of magnitude (Fig. 5A, inset; Table S5). The corresponding dissociation rates were also lowered significantly, suggesting that EGCG may act by simply slowing down aggregation rather than destabilizing misfolded oligomers.

**Figure 5:**
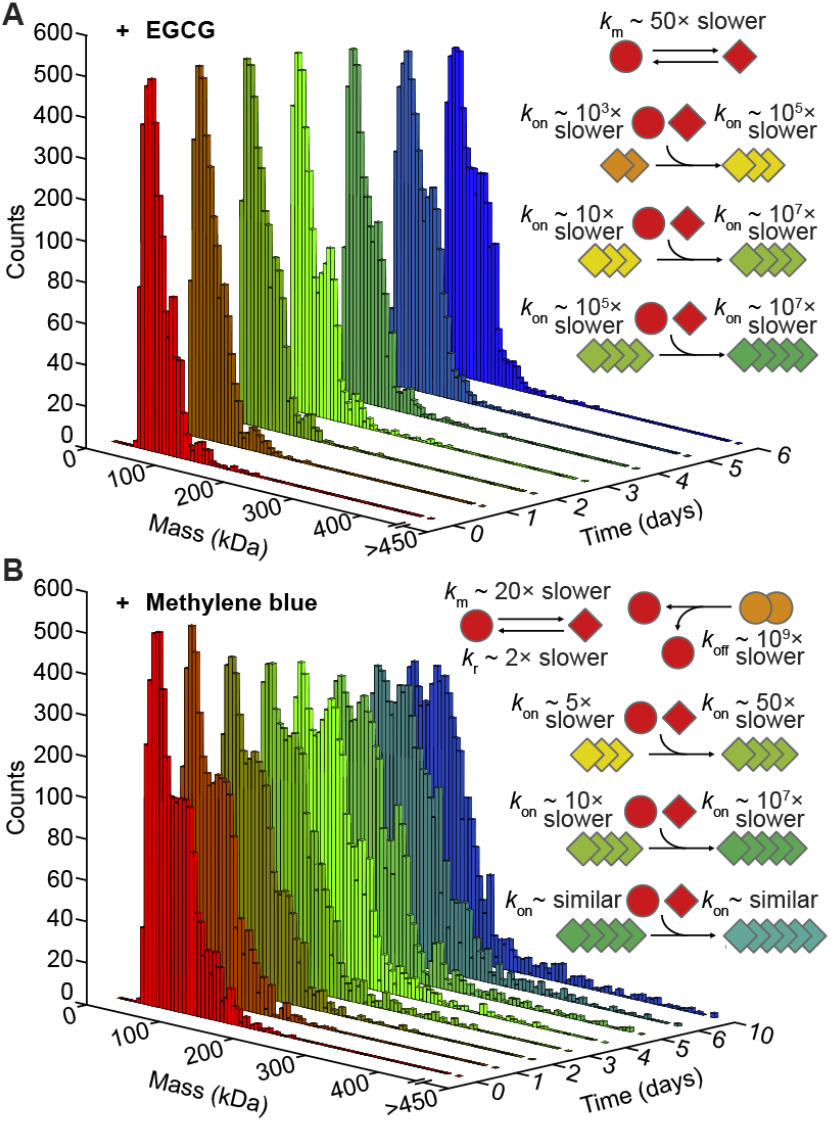
Effects of aggregation inhibitors. (A) EGCG effectively arrests aggregation by greatly slowing growth of all misfolded oligomers. (B) Methylene blue greatly slows but does not stop aggregation, by inhibiting monomer misfolding and creating a bottleneck at tetramers. Insets: key changes in rates.

Turning to methylene blue, we found that here (unlike with EGCG) a distinct tetramer peak was observed after 24 hours, indicating that aggregation is occurring (Fig. 5B). However, even though scattered higher-order oligomers were observed at later times, no well-defined peaks could be observed, suggesting that the tetramer may act as a choke point in the aggregation mechanism. Kinetic modeling of the data (Fig S5) confirmed this picture, yielding a 10 million-fold reduction in the rate of misfolded monomer addition to the tetramer, but suggested that addition and removal rates for the pentamer and larger were not much affected (Fig. 5B, inset; Table S6). In addition, methylene blue stabilized the native form of the monomer.

Taken together, these results imply a mechanism of action distinct from EGCG, stabilizing misfolded tetramers of tau but failing to slow subsequent aggregation steps, leading to the slow accumulation of larger oligomers.

## DISCUSSION

Mapping the discrete steps in the aggregation cascade to define aggregation mechanisms and pinpoint how inhibitors act on them is critical for understanding aggregation as a biophysical process, defining therapeutic targets in the cascade, and guiding discovery and development of therapeutic candidates. By giving direct access to the time evolution of all small-oligomer populations, SMMP allows the mechanisms of aggregation and effects of inhibitors to be probed at an unprecedented level of detail: since the growth/decay curves for every oligomeric species observed are coupled, the microscopic rates for each step in the cascade can be obtained from global modeling without overfitting. Furthermore, direct detection of individual unlabeled molecules avoids interpretative difficulties that can arise from labels that may alter the behavior of the protein (whether overtly or subtly) or from detection methods with biased size sensitivity.

Comparing the results of SMMP and ThT fluorescence measurements of P301L tau aggregation is particularly instructive regarding the powerful insights into aggregation mechanisms offered by SMMP. Our analysis of ThT fluorescence using current best practices implies that P301L tau aggregates via saturating elongation with either fragmentation or secondary nucleation. Yet such a mechanism is grossly inconsistent with the oligomer populations observed directly by SMMP. Indeed, it turns out that a simple nucleation-elongation model matches the observed oligomer dynamics better (Fig. S3) even though it fits the ThT data poorly. These results show that ThT fluorescence may not be reliable for inferring aggregation mechanisms: not only did the standard models for fitting fibril kinetics inadequately capture the oligomer dynamics, but improving the ThT fits actually led to a worse description of the oligomer populations. In particular, adding secondary processes to optimize fitting of long lag phases followed by rapid growth led to overestimation of the prevalence of the smallest oligomers and underestimation of that of larger oligomers (Fig. 4). In contrast, fitting the oligomer dynamics observed by SMMP also provided a reasonable prediction of the fibril kinetics observed by ThT fluorescence. These results imply that SMMP on its own captures the essence of the aggregation mechanism and that combining both methods by fitting SMMP and ThT data simultaneously yields a model that accurately describes the dynamics across the full range of the cascade, from early oligomers to fibrils (Fig. 3). By showing that secondary processes are not needed to explain aggregation here, these results also contradict claims that such processes are quasiuniversal in self-replicating aggregation [26].

Turning to the aggregation mechanism deduced here for P301L tau, we find that it is similar to that seen for WT tau [17], featuring two pathways: one off-pathway to aggregation, involving a native monomer-dimer-trimer equilibrium, and the other on-pathway to large aggregates, with the two connected via a conformational change in the monomer. In contrast to WT tau, however, here the monomer misfolding occurs spontaneously instead of being induced externally. A second major difference is that oligomer growth is much slower, occurring over days rather than minutes, owing to much slower monomer addition rates. Most intriguingly, whereas WT tau oligomer growth suddenly accelerated and became nearly irreversible after oligomers grew to pentamers, dividing the kinetics into a slow nucleation-like phase (featuring reversible transitions leading to pentamers) followed by rapid growth, no such division was seen for P301L tau. Instead, the addition/removal rates changed more gradually, and no nucleation-like phase prior to accelerated growth could be discerned. This result runs counter to the notion that aggregation is a nucleated process in which the formation of an unstable oligomeric species acts as an energetic barrier to subsequent rapid growth [37]. Although lag phases in aggregation are typically explained as a consequence of the kinetic barrier created by a critical nucleus, here we see that multiple reversible kinetic steps can combine to generate a lag phase in the absence of a critical nucleus. This finding highlights the importance of directly observing the steps that are proposed to play critical mechanistic roles, to avoid assumptions that may lead to incorrect interpretations.

The ability of SMMP to deduce microscopic rates in each step of the aggregation mechanism is particularly powerful for studying aggregation inhibitors, as it enables identification of which steps in the cascade are affected. In the case of EGCG and methylene blue, both had similar effects on fibrillization, strongly suppressing it, and also broadly similar effects on oligomers, although methylene blue did allow the very slow appearance of oligomers larger than tetramers. However, the kinetic analysis of oligomers revealed important mechanistic differences, suggesting that methylene blue acts primarily as a bottleneck to tetramer growth, in contrast to the more broad-spectrum inhibition of EGCG across all oligomer sizes. This effect may account for the failure of methylene blue to protect against tau toxicity, as it could establish a persistent population of toxic small oligomers as suggested previously [24]. As for EGCG, the global slowing of all aggregation suggests that it may bind promiscuously to several different oligomeric species, an idea that is supported by its ability to bind to a wide range of proteins [23]. The ability to distinguish the mechanistic effects of different compounds by assessing their impact on each step of the cascade makes SMMP ideally suited for developing and characterizing protein aggregation inhibitors.

Although SMMP is a powerful tool for studying aggregation, it does have limitations. The most important is the detection limit for light scattering: molecules with low molecular weight do not scatter sufficient light to be detected [12]. Practical detection limits of ∼20–30 kDa pose a challenge for applying SMMP to study a number of aggregation-prone proteins and peptides of biomedical relevance (including aβ peptide, α-synuclein, islet amyloid polypeptide, and others). The need to convert postulated models into systems of differential equations for numerical integration during fitting can also be challenging, but this approach offers compensating advantages: simplistic assumptions can be avoided (*e*.*g*. size-independent rates and coupling of nucleation and ThT-positivity, as required in analytical models [25]), and additional process can be easily added (*e*.*g*. addition/removal of oligomers, secondary growth processes), allowing for a more complete and realistic modeling of mechanisms. The ability of SMMP to determine protein aggregation mechanisms, bridging the gap between early oligomers and late fibrils, positions SMMP as a valuable tool for protein aggregation research and pharmaceutical development.

## METHODS

### Sample preparation

The 0N4R isoform of the tau mutant P301L was purified as described previously for WT tau [17]. All buffers used for tau dilution or measurement were double filtered through 0.22-μm syringe filters to eliminate contaminants that could be detectable with SMMP.

### Tau aggregation

For both SMMP and ThT fluorescence measurements, P301L tau was suspended at the desired final concentration in phosphate buffered saline (PBS), pH 7.5, containing 2 mM DTT and 0.1 mM PMSF. To preserve a reducing environment over the course of the experiments, DTT was added every 24 hr to the incubation at a final additional concentration of 1 mM. Samples were incubated at 37 °C while shaking continuously at 300 rpm, with aliquots being removed and placed on ice at each time-point for measurement. For SMMP assays of inhibitors, a final concentration of 50 μM inhibitor was added at the start of the incubation.

### SMMP measurements

SMMP measurements were taken with the OneMP mass photometer (Refeyn), performed as done previously for WT tau [17]. Briefly, the instrument was calibrated with bovine serum albumin (BSA) and apoferritin using manufacturer protocols. Measurements were done using protein samples incubated at 10 μM; inhibitor, if present, was added to a final concentration of 50 μM. At each time-point, samples were diluted to a final tau concentration of 40 nM in the measuring well: first they were diluted serially to 120 nM in PBS outside the well, then 10 μL of PBS was added to the well for a 1-min measurement of blank buffer to ensure buffer cleanliness, and lastly 5 μL of the diluted protein sample was added to the well to achieve the desired final concentration for measurement of oligomer populations. Movies of the scattering were recorded for 1 min and analyzed as described below. Measurements for each time-point and experimental condition (inhibitor/no inhibitor) were replicated three times using different aggregation reactions.

### ThT fluorescence measurements

Aliquots of tau incubated as described above at the desired concentrations (2.5, 5, 7.5, 10, 15, and 20 μM) were taken every 12 hours and thioflavin-T (ThT) was added to a final concentration of 15 μM. ThT fluorescence was measured from 96-well plates in a microplate reader, with 150 μL well volume, exciting at 440 nm excitation and measuring emission at 485 nm (20-nm bandwidth in each case). Plates were shaken for 30 s (300 rpm double orbital shaking) before measurement.

### SMMP data analysis

Movies of the scattering were analyzed using the DiscoverMP software (Refeyn) as done previously for WT tau [17]. Histograms of binding events for masses between 0 and 1500.2 kDa were generated with a bin-width of 5.2 kDa. Peaks in the mass spectrum *P*(*m*) were fit to Gaussians of constant width spaced equally at integer multiples of the mass of tau. For measurements of P301L tau alone, the mass used in the fits was determined from fitting the spectrum of the 96-hr time-point (which had the most peaks) to the sum of 10 Gaussian peaks multiplied by a sigmoid function reflecting the resolution limit of the instrument:

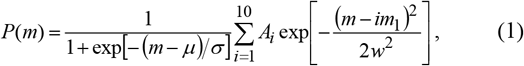

where *A*_*i*_ is the amplitude of the peak for *i*-mers, *m*_1_ the monomer mass, *w* the peak width, *μ* the sigmoid centre, and *σ* the sigmoid width. The fit result *m*_1_ = 43.45 kDa from the 96-hr time-point was then used as a constraint when fitting all time-points to Eq. 1. Counts for monomers through 10-mers were found from the area under the peak divided by the bin width. To correct for the effect of diffusional undercounting of larger species, we multiplied each of these counts by *n*^1/6^ (for *n* = oligomer size) as done previously for WT tau [17]. The counts for 11to 15-mers and 16- to 25-mers were found by summing the diffusion-corrected counts between 452.4 and 670.8 kDa or 670.8 and 1097.2 kDa, respectively. For measurements in the presence of inhibitors, the number of peaks included in the fitting differed: 3 for EGCG, 10 for methylene blue. Data from day 3 was used to find *m*_1_ with EGCG present (*m*_1_ = 43.78 kDa), and from day 5 with methylene blue present (*m*_1_ = 43.52 kDa). With EGCG present, the counts for 4- to 10-mers were combined by summing the diffusion-corrected counts between 150.8 and 462.8 kDa. All histogram fitting was done with Igor Pro 8.0.4.

Kinetic models were implemented as systems of coupled first-order differential equations and solved using the ode45 function in MATLAB R2021a (MathWorks, 2021). Equations for both models used are given in Supporting Information. Model parameters were found by random initialization followed by log-space gradient descent optimization with the trust-region-reflective algorithm in MATLAB R2021a. Once converged, we then performed local sampling to avoid shallow local minima, followed by further gradient descent if a locally better parameter set was found. This procedure was run for up to 100 million starting points per kinetic model. Standard deviations of the fit parameters were first estimated crudely by reducing the value of each parameter independently until fit χ^2^ increased by 1. We then randomly sampled the parameter space using a normal distribution centered on the optimum with these estimated standard deviations: by assigning a *p*- value of exp(−χ^2^/2) to each parameter set, we estimated the parameter covariance matrix by performing this sampling 50,000 times per model, reporting parameter errors as the square root of the diagonal matrix entries. Rates were converted to molar units using the count/molarity conversion factor found by dividing the total number of monomers units observed at time zero by the actual monomer concentration (10 μM). Since the oligomers in these models can grow to arbitrary size, for computational feasibility we treated 100-mers as a sink term (where monomers can be added to but not removed); none of the models showed any counts in the sink term, implying that the sink term did not affect the results.

### ThT data analysis

ThT data were analyzed by direct numerical integration of the models described for the SMMP analysis, but using 1000-mers as the sink term. The contribution to the ThT signal from oligomers of size *i* was calculated by multiplying the number of monomer unit counts from that oligomer (*i*.*e. i×N*_*i*_) by the sigmoid function [1 + exp[−(*i* − *i*_ThT_)/*α*)] ^−1^, where *i*_ThT_ is the oligomer size at which aggregates become ThT-sensitive and *α* controls the width of the transition between ThT sensitivity and - insensitivity. These values were then summed over all oligomer populations to get the total ThT signal at that timepoint. For the ThT forecasting (Fig. 3C, dashed line), *i*_ThT_ and α were fit only to the ThT data while keeping the other kinetic parameters constant, using the same fitting strategy employed for the SMMP analysis. For the ThT fit (Fig. 3C, solid line) we assumed that the on and off-rates changed smoothly along the sigmoid from the values for 16- to 25-mers to the rates describing growth/decay of ThT-positive fibrils (see Supporting Information). We then performed a global fit of all parameters to both the SMMP and ThT data; the small oligomer rates changed very little from the fit to the SMMP data alone, leading to nearly identical predictions for the dynamics of monomers through 25-mers (Fig. S6).

Global fitting of concentration-dependent ThT data was performed using the online platform Amylofit [25]. We fitted the data to all unseeded models available on the platform, using a fixed primary and secondary nucleation order of 2. Fit quality was assessed using the root mean-squared deviation (RMSD) value. To assess how well fits predicted our SMMP data, we converted the models with the lowest and highest RMSD values (respectively, saturating elongation & fragmentation and nucleation & elongation) into equivalent systems of first-order differential equations that allowed us to track all oligomer populations separately (see Supporting Information). The differential forms of the Amylofit models were used to ensure that these conversions captured the corresponding analytical models correctly. Since the elongation rate is not uniquely determined from fits of unseeded models, we used the Amylofit results as constraints to perform a single-parameter fit of both converted models to the data for 1- to 25-mers.

## Acknowledgements

This work was funded by the Natural Sciences and Engineering Research Council of Canada (grant numbers RTI-2020-00301, RGPIN-2018-04673, and ALLRP 586226-23 to MTW; grant number RGPIN-2019- 06230 to SAM).

## Author contributions

AL, SSP, SAM, and MTW designed research; SSP and AY prepared samples; SSP and GH collected data; AL and JEM developed analysis methods; AL and JEM analyzed data; AL and MTW wrote manuscript; AL, SSP, SAM, and MTW edited manuscript.

## Supporting Information

### Equations for SMMP kinetic modeling

The differential equations describing the aggregation models used are described here.

#### Model 1 (Fig. S1): single pathway

This model involves a single pathway with addition/removal of native monomers to oligomers of all sizes. In the equations below, *n*_*i*_ is the number of oligomers of size *i*, and the various rates *k* describe addition of a native monomer (superscript +n) or removal of one (superscript −n) onto/from an oligomer of size *i* (subscript n_*i*_). Oligomers of size *N* = 100 are taken to be a sink term (*i*.*e*., monomers can be added to them but never removed).

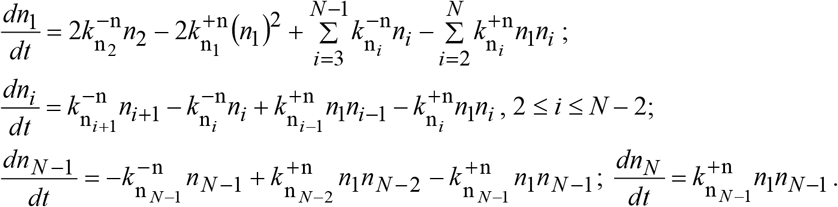

#### Model 2 (Fig. 3A): monomer misfolding

Here, native monomers can spontaneously convert to a misfolded form which can grow into misfolded oligomers by addition of either native or misfolded monomers. Native oligomers (denoted *n*_*i*_ as above) exist in a monomer-dimer-trimer equilibrium, whereas misfolded oligomers (where the number of oligomers of size *i* is *m*_*i*_) can grow without bound. Oligomers of size *N* = 100 are taken to be a sink term. Rates (*k*) describe addition of a native monomer (superscript +n), addition of a misfolded monomer (superscript +m) or removal of a misfolded monomer (superscript −m) from a native or misfolded oligomer of size *i* (subscript m_*i*_). *k*_m_ and *k*_r_ describe the misfolding and refolding rates of the monomeric species.

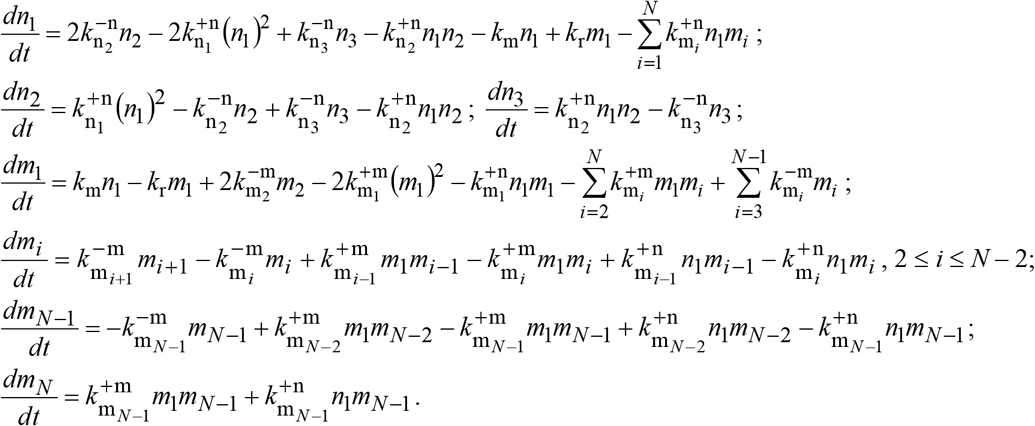

Rates of native monomer removal from native oligomers were calculated assuming an initial monomerdimer-trimer equilibrium using the corresponding addition rates and the initial oligomer counts:

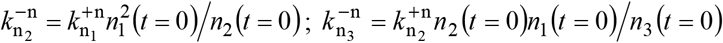

#### Model 3 (Fig. 3C, dashed line): monomer misfolding with ThT forecasting

This model is identical to Model 2, but increases the sink term to *N =* 1000 to allow for large, ThT-positive fibrils. Aggregates are assumed to become ThT positive in a size-dependent fashion, following a sigmoid with center *i*_ThT_ and width α. The ThT signal, *M*(*t*), is then calculated by summing up the contribution of all monomers in these ThT-positive aggregates:

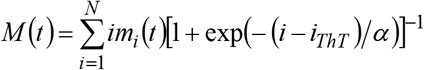

To fit the normalized ThT data, this function was normalized by dividing by its value at the final timepoint.

#### Model 4 (Fig. 3C, solid line): monomer misfolding with ThT-positive rate switching

This model is equivalent to model 3 for oligomers smaller than 25-mers (the largest oligomer monitored by SMMP), but includes different addition/removal rates for larger, ThT-positive oligomers. Rates are assumed to transition smoothly from the values for 25-mers to the values for ThT-positive oligomers following the same sigmoid function assumed for ThT-positivity. Hence for *i* > 25, addition/removal rates gradually transition to their ThT-positive values depending on the sigmoid center *i*_ThT_ and width *α*:

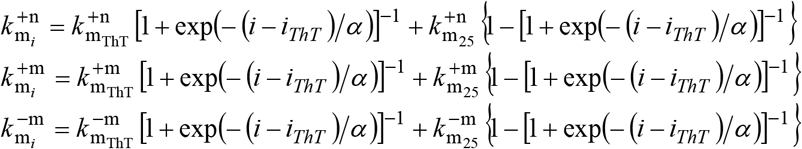

### Equations for ThT fluorescence converted from analytical models

The analytical models for unseeded nucleation-elongation and saturating elongation-fragmentation from Ref. [1], which were used to fit the global ThT kinetics, were converted into differential equations describing the populations of all oligomers. We assumed an entirely monomeric initial condition for both models, an assumption also made by the original equations.

#### Nucleation-elongation model (Fig. S3)

This model is similar to Model 1 described above, but with a separate nucleation step for dimer formation and no dissociation rates.

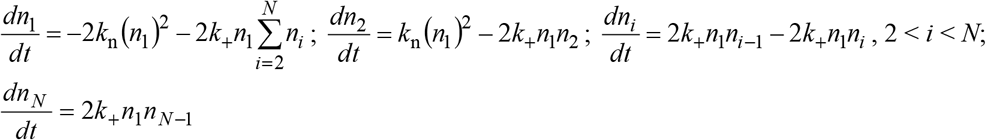

Here, *k*_+_ is the elongation rate, *k*_n_ is the nucleation rate, and *N* = 500 is the sink term. The factor of two in front of the elongation rates comes from the original model for fibrillization and is used here to ensure parameters are consistent across both models. The elongation rate is not uniquely specified from the ThT fit, hence we treated it as a free parameter constraining *k*_n_. The fit to the fibril model in Amylofit yielded *k*_+_*k*_n_ = 1.68 × 10^8^ M^−2^ days^−2^, which was used as a constraint when fitting the other parameters. The resulting one-parameter fit gave *k*_+_ = 1.03 × 10^5^ M^−1^ days^−1^ and *k*_n_ = 1.64 × 10^3^ M^−1^ days^−1^.

#### Saturating elongation-fragmentation (Fig. 4C, Fig. S2)

In addition to a saturating growth rate, this model assumes that dimers and larger can spontaneously break into any combination of their constituent components. Following Ref. [2], we assume that oligomers are linear polymers and can fragment with equal probability at any link between monomers, leading to the fragmentation rate being approximated as proportional to the total aggregated monomer concentration (the assumption made in the Amylofit models).

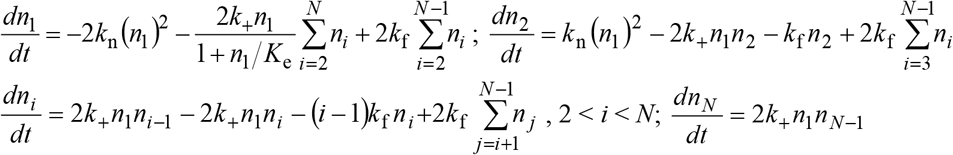

Here, *k*_f_ is the fragmentation rate, *K*_e_ is the saturating elongation concentration, and all other parameters are defined as in the nucleation-elongation model. The sink term is unable to fragment, but the sink size of *N* = 500 was chosen such that no oligomers of the sink size were formed within the timescale of the numerical integration. Again the elongation rate is not uniquely specified from the ThT fit, hence we treated it as a free parameter constraining *k*_n_ and *k*_f_. Amylofit yielded *k*_+_*k*_n_ = 2.73 × 10^8^ M^−2^ days^−2^, *k*_+_*k*_f_ = 3.06 × 10^3^ M^−1^ days^−2^, and *K*_e_ = 2.04 × 10^−6^ M, which were used as fitting constraints. The resulting one-parameter fit gave *k*_+_ = 5.92 × 10^4^ M^−1^ days^−1^, *k*_n_ = 4.62 × 10^3^ M^−1^ days^−1^, and *k*_f_ = 5.17 × 10^−2^ days^−1^.

**Table S1:**
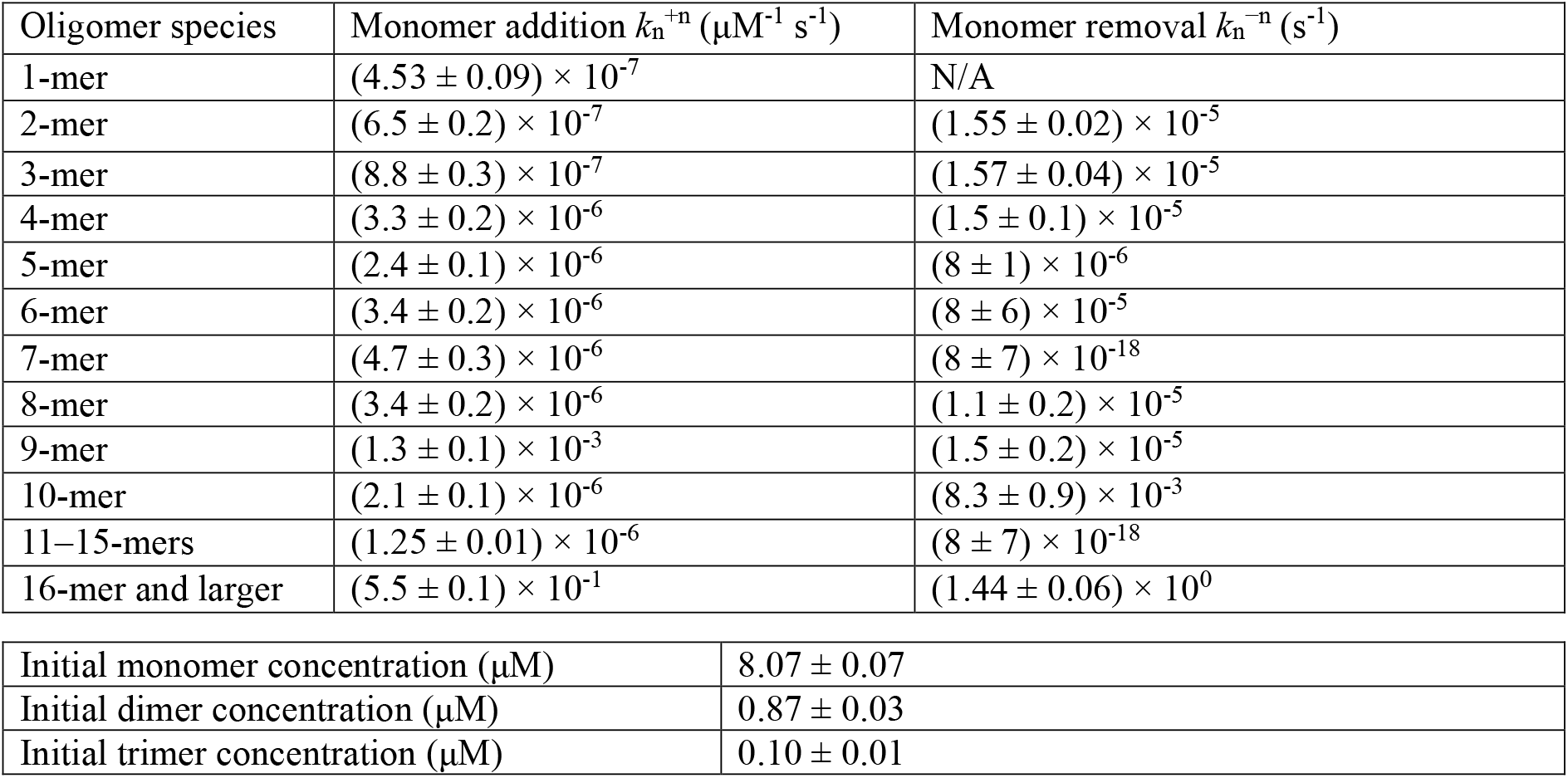
SMMP fit results—model 1. Results from fitting SMMP data in Fig 3 (no inhibitor) to model 1.

| Oligomer species | Monomer addition $k_n^{+n} (\mu\text{M}^{-1} \text{ s}^{-1})$ | Monomer removal $k_n^{-n} (\text{s}^{-1})$ |
| --- | --- | --- |
| 1-mer | $(4.53 \pm 0.09) \times 10^{-7}$ | N/A |
| 2-mer | $(6.5 \pm 0.2) \times 10^{-7}$ | $(1.55 \pm 0.02) \times 10^{-5}$ |
| 3-mer | $(8.8 \pm 0.3) \times 10^{-7}$ | $(1.57 \pm 0.04) \times 10^{-5}$ |
| 4-mer | $(3.3 \pm 0.2) \times 10^{-6}$ | $(1.5 \pm 0.1) \times 10^{-5}$ |
| 5-mer | $(2.4 \pm 0.1) \times 10^{-6}$ | $(8 \pm 1) \times 10^{-6}$ |
| 6-mer | $(3.4 \pm 0.2) \times 10^{-6}$ | $(8 \pm 6) \times 10^{-5}$ |
| 7-mer | $(4.7 \pm 0.3) \times 10^{-6}$ | $(8 \pm 7) \times 10^{-18}$ |
| 8-mer | $(3.4 \pm 0.2) \times 10^{-6}$ | $(1.1 \pm 0.2) \times 10^{-5}$ |
| 9-mer | $(1.3 \pm 0.1) \times 10^{-3}$ | $(1.5 \pm 0.2) \times 10^{-5}$ |
| 10-mer | $(2.1 \pm 0.1) \times 10^{-6}$ | $(8.3 \pm 0.9) \times 10^{-3}$ |
| 11–15-mers | $(1.25 \pm 0.01) \times 10^{-6}$ | $(8 \pm 7) \times 10^{-18}$ |
| 16-mer and larger | $(5.5 \pm 0.1) \times 10^{-1}$ | $(1.44 \pm 0.06) \times 10^0$ |

|  |  |
| --- | --- |
| Initial monomer concentration ($\mu\text{M}$) | $8.07 \pm 0.07$ |
| Initial dimer concentration ($\mu\text{M}$) | $0.87 \pm 0.03$ |
| Initial trimer concentration ($\mu\text{M}$) | $0.10 \pm 0.01$ |

**Table S2:**
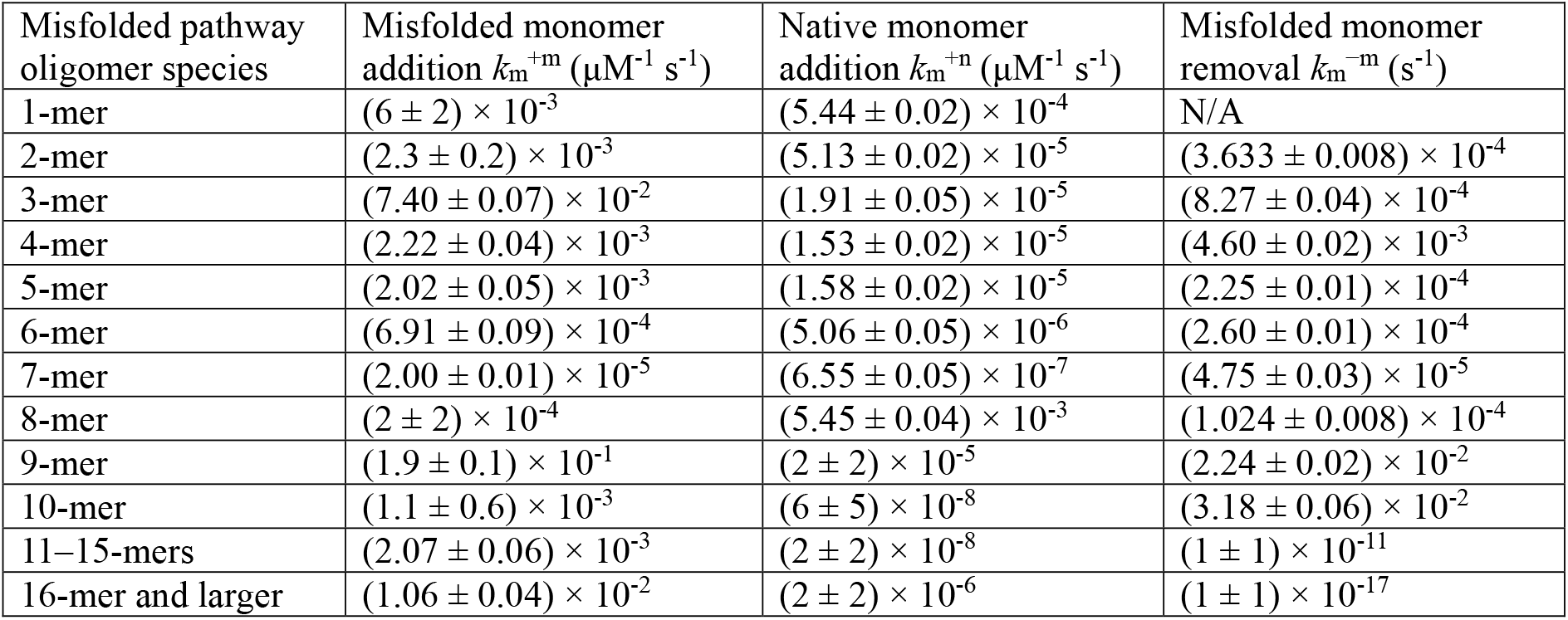
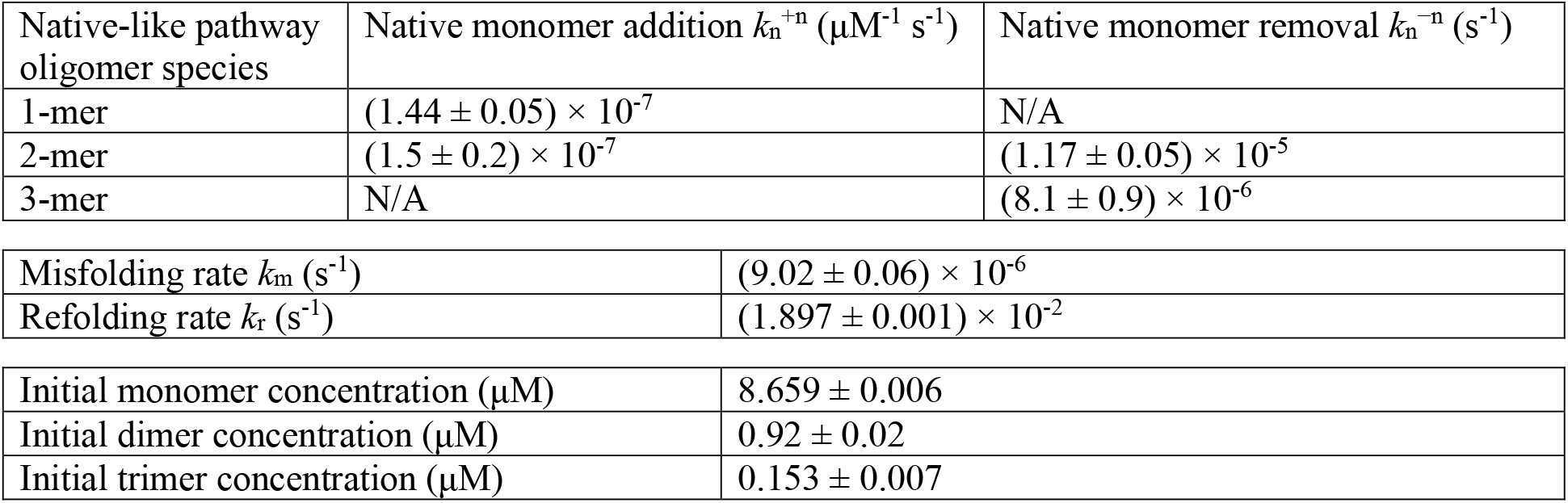
SMMP fit results—model 2. Results from fitting SMMP data in Fig. 3 (no inhibitor) to model 2.

| Misfolded pathway oligomer species | Misfolded monomer addition $k_m^{+m} (\mu\text{M}^{-1} \text{ s}^{-1})$ | Native monomer addition $k_m^{+n} (\mu\text{M}^{-1} \text{ s}^{-1})$ | Misfolded monomer removal $k_m^{-m} (\text{s}^{-1})$ |
| --- | --- | --- | --- |
| 1-mer | $(6 \pm 2) \times 10^{-3}$ | $(5.44 \pm 0.02) \times 10^{-4}$ | N/A |
| 2-mer | $(2.3 \pm 0.2) \times 10^{-3}$ | $(5.13 \pm 0.02) \times 10^{-5}$ | $(3.633 \pm 0.008) \times 10^{-4}$ |
| 3-mer | $(7.40 \pm 0.07) \times 10^{-2}$ | $(1.91 \pm 0.05) \times 10^{-5}$ | $(8.27 \pm 0.04) \times 10^{-4}$ |
| 4-mer | $(2.22 \pm 0.04) \times 10^{-3}$ | $(1.53 \pm 0.02) \times 10^{-5}$ | $(4.60 \pm 0.02) \times 10^{-3}$ |
| 5-mer | $(2.02 \pm 0.05) \times 10^{-3}$ | $(1.58 \pm 0.02) \times 10^{-5}$ | $(2.25 \pm 0.01) \times 10^{-4}$ |
| 6-mer | $(6.91 \pm 0.09) \times 10^{-4}$ | $(5.06 \pm 0.05) \times 10^{-6}$ | $(2.60 \pm 0.01) \times 10^{-4}$ |
| 7-mer | $(2.00 \pm 0.01) \times 10^{-5}$ | $(6.55 \pm 0.05) \times 10^{-7}$ | $(4.75 \pm 0.03) \times 10^{-5}$ |
| 8-mer | $(2 \pm 2) \times 10^{-4}$ | $(5.45 \pm 0.04) \times 10^{-3}$ | $(1.024 \pm 0.008) \times 10^{-4}$ |
| 9-mer | $(1.9 \pm 0.1) \times 10^{-1}$ | $(2 \pm 2) \times 10^{-5}$ | $(2.24 \pm 0.02) \times 10^{-2}$ |
| 10-mer | $(1.1 \pm 0.6) \times 10^{-3}$ | $(6 \pm 5) \times 10^{-8}$ | $(3.18 \pm 0.06) \times 10^{-2}$ |
| 11–15-mers | $(2.07 \pm 0.06) \times 10^{-3}$ | $(2 \pm 2) \times 10^{-8}$ | $(1 \pm 1) \times 10^{-11}$ |
| 16-mer and larger | $(1.06 \pm 0.04) \times 10^{-2}$ | $(2 \pm 2) \times 10^{-6}$ | $(1 \pm 1) \times 10^{-17}$ |

| Native-like pathway oligomer species | Native monomer addition $k_n^{+n}$ ($\mu\text{M}^{-1} \text{s}^{-1}$) | Native monomer removal $k_n^{-n}$ ($\text{s}^{-1}$) |
| --- | --- | --- |
| 1-mer | $(1.44 \pm 0.05) \times 10^{-7}$ | N/A |
| 2-mer | $(1.5 \pm 0.2) \times 10^{-7}$ | $(1.17 \pm 0.05) \times 10^{-5}$ |
| 3-mer | N/A | $(8.1 \pm 0.9) \times 10^{-6}$ |

|  |  |
| --- | --- |
| Misfolding rate $k_m$ ($\text{s}^{-1}$) | $(9.02 \pm 0.06) \times 10^{-6}$ |
| Refolding rate $k_r$ ($\text{s}^{-1}$) | $(1.897 \pm 0.001) \times 10^{-2}$ |
| Initial monomer concentration ($\mu\text{M}$) | $8.659 \pm 0.006$ |
| Initial dimer concentration ($\mu\text{M}$) | $0.92 \pm 0.02$ |
| Initial trimer concentration ($\mu\text{M}$) | $0.153 \pm 0.007$ |

**Table S3:** ThT fit results—model 3. Results from fitting ThT data in Fig 3 (no inhibitor) to model 3.

|  |  |
| --- | --- |
| ThT-positive sigmoid center size $i_{\text{ThT}}$ | $89 \pm 3$ |
| ThT-positive sigmoid width $\alpha$ | $(5 \pm 6) \times 10^{-5}$ |

**Table S4:**
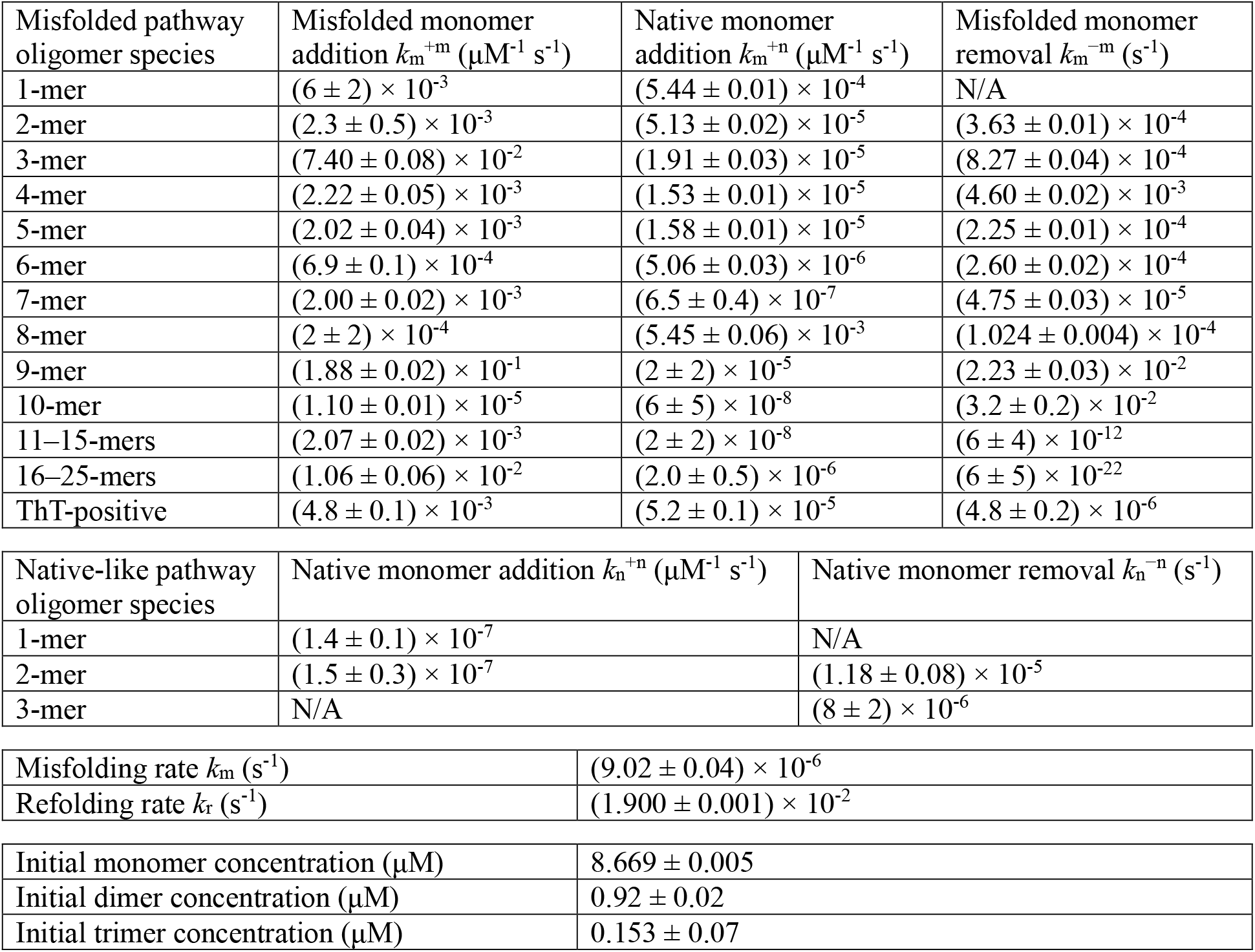
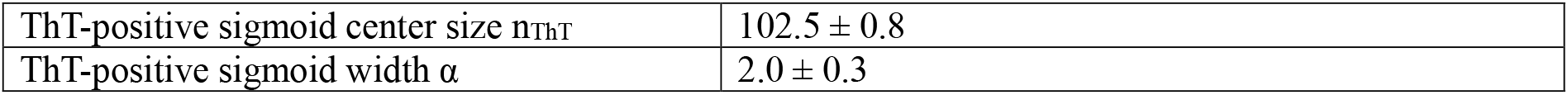
Combined SMMP and ThT fit results—model 4. Results from simultaneous fitting of SMMP and ThT data in Fig 3 (no inhibitor) to model 4.

| Misfolded pathway oligomer species | Misfolded monomer addition $k_m^{+m}$ ($\mu\text{M}^{-1} \text{s}^{-1}$) | Native monomer addition $k_m^{+n}$ ($\mu\text{M}^{-1} \text{s}^{-1}$) | Misfolded monomer removal $k_m^{-m}$ ($\text{s}^{-1}$) |
| --- | --- | --- | --- |
| 1-mer | $(6 \pm 2) \times 10^{-3}$ | $(5.44 \pm 0.01) \times 10^{-4}$ | N/A |
| 2-mer | $(2.3 \pm 0.5) \times 10^{-3}$ | $(5.13 \pm 0.02) \times 10^{-5}$ | $(3.63 \pm 0.01) \times 10^{-4}$ |
| 3-mer | $(7.40 \pm 0.08) \times 10^{-2}$ | $(1.91 \pm 0.03) \times 10^{-5}$ | $(8.27 \pm 0.04) \times 10^{-4}$ |
| 4-mer | $(2.22 \pm 0.05) \times 10^{-3}$ | $(1.53 \pm 0.01) \times 10^{-5}$ | $(4.60 \pm 0.02) \times 10^{-3}$ |
| 5-mer | $(2.02 \pm 0.04) \times 10^{-3}$ | $(1.58 \pm 0.01) \times 10^{-5}$ | $(2.25 \pm 0.01) \times 10^{-4}$ |
| 6-mer | $(6.9 \pm 0.1) \times 10^{-4}$ | $(5.06 \pm 0.03) \times 10^{-6}$ | $(2.60 \pm 0.02) \times 10^{-4}$ |
| 7-mer | $(2.00 \pm 0.02) \times 10^{-3}$ | $(6.5 \pm 0.4) \times 10^{-7}$ | $(4.75 \pm 0.03) \times 10^{-5}$ |
| 8-mer | $(2 \pm 2) \times 10^{-4}$ | $(5.45 \pm 0.06) \times 10^{-3}$ | $(1.024 \pm 0.004) \times 10^{-4}$ |
| 9-mer | $(1.88 \pm 0.02) \times 10^{-1}$ | $(2 \pm 2) \times 10^{-5}$ | $(2.23 \pm 0.03) \times 10^{-2}$ |
| 10-mer | $(1.10 \pm 0.01) \times 10^{-5}$ | $(6 \pm 5) \times 10^{-8}$ | $(3.2 \pm 0.2) \times 10^{-2}$ |
| 11–15-mers | $(2.07 \pm 0.02) \times 10^{-3}$ | $(2 \pm 2) \times 10^{-8}$ | $(6 \pm 4) \times 10^{-12}$ |
| 16–25-mers | $(1.06 \pm 0.06) \times 10^{-2}$ | $(2.0 \pm 0.5) \times 10^{-6}$ | $(6 \pm 5) \times 10^{-22}$ |
| ThT-positive | $(4.8 \pm 0.1) \times 10^{-3}$ | $(5.2 \pm 0.1) \times 10^{-5}$ | $(4.8 \pm 0.2) \times 10^{-6}$ |

| Native-like pathway oligomer species | Native monomer addition $k_n^{+n}$ ($\mu\text{M}^{-1} \text{s}^{-1}$) | Native monomer removal $k_n^{-n}$ ($\text{s}^{-1}$) |
| --- | --- | --- |
| 1-mer | $(1.4 \pm 0.1) \times 10^{-7}$ | N/A |
| 2-mer | $(1.5 \pm 0.3) \times 10^{-7}$ | $(1.18 \pm 0.08) \times 10^{-5}$ |
| 3-mer | N/A | $(8 \pm 2) \times 10^{-6}$ |

|  |  |
| --- | --- |
| Misfolding rate $k_m$ ($\text{s}^{-1}$) | $(9.02 \pm 0.04) \times 10^{-6}$ |
| Refolding rate $k_r$ ($\text{s}^{-1}$) | $(1.900 \pm 0.001) \times 10^{-2}$ |
| Initial monomer concentration ($\mu\text{M}$) | $8.669 \pm 0.005$ |
| Initial dimer concentration ($\mu\text{M}$) | $0.92 \pm 0.02$ |
| Initial trimer concentration ($\mu\text{M}$) | $0.153 \pm 0.07$ |
| ThT-positive sigmoid center size $n_{\text{ThT}}$ | $102.5 \pm 0.8$ |
| ThT-positive sigmoid width $\alpha$ | $2.0 \pm 0.3$ |

**Table S5:**
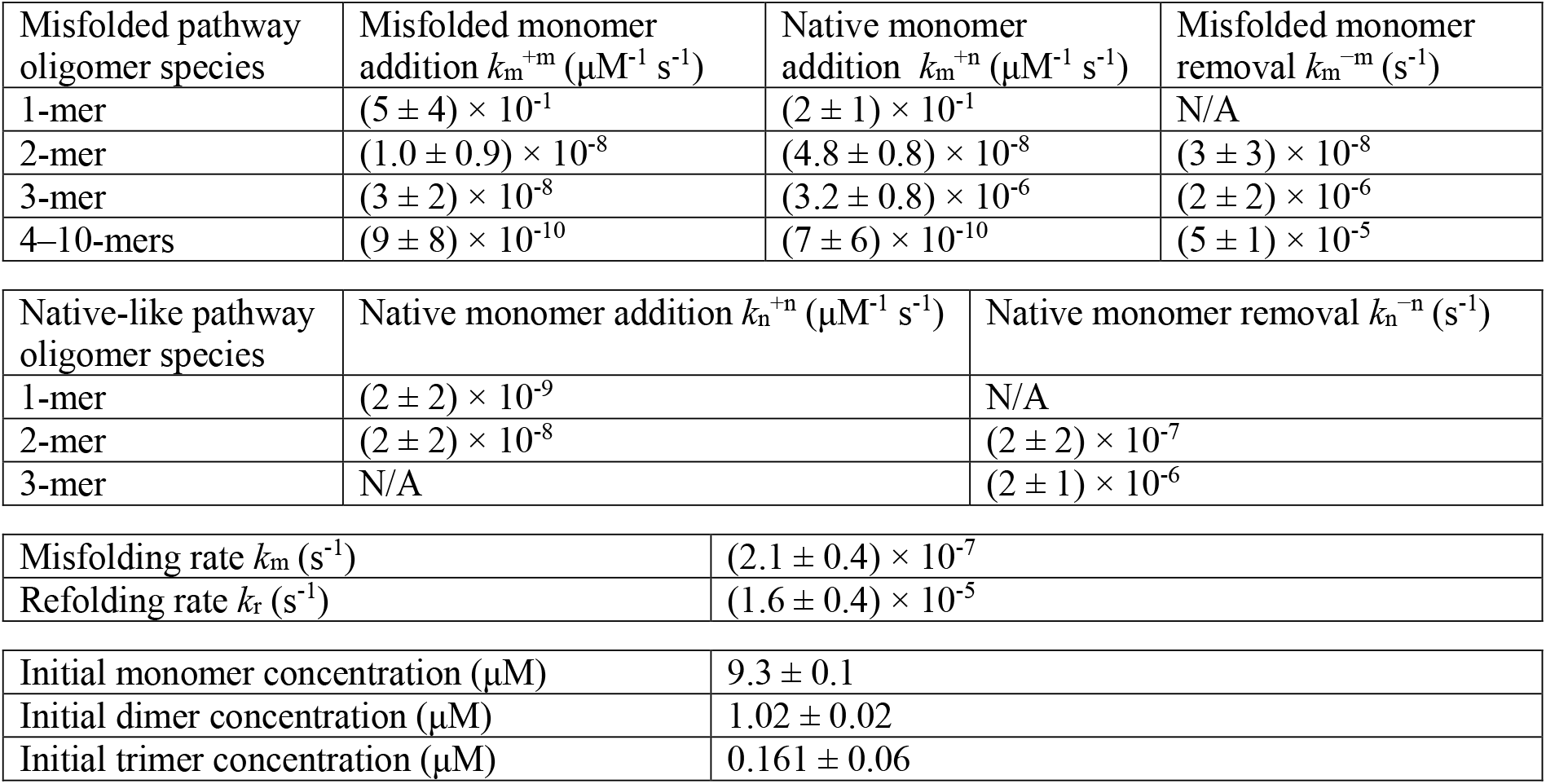
SMMP fit results with EGCG. Results from fitting SMMP data in Fig. 5A to model 2.

| Misfolded pathway oligomer species | Misfolded monomer addition $k_m^{+m}$ ($\mu\text{M}^{-1} \text{s}^{-1}$) | Native monomer addition $k_m^{+n}$ ($\mu\text{M}^{-1} \text{s}^{-1}$) | Misfolded monomer removal $k_m^{-m}$ ($\text{s}^{-1}$) |
| --- | --- | --- | --- |
| 1-mer | $(5 \pm 4) \times 10^{-1}$ | $(2 \pm 1) \times 10^{-1}$ | N/A |
| 2-mer | $(1.0 \pm 0.9) \times 10^{-8}$ | $(4.8 \pm 0.8) \times 10^{-8}$ | $(3 \pm 3) \times 10^{-8}$ |
| 3-mer | $(3 \pm 2) \times 10^{-8}$ | $(3.2 \pm 0.8) \times 10^{-6}$ | $(2 \pm 2) \times 10^{-6}$ |
| 4–10-mers | $(9 \pm 8) \times 10^{-10}$ | $(7 \pm 6) \times 10^{-10}$ | $(5 \pm 1) \times 10^{-5}$ |

| Native-like pathway oligomer species | Native monomer addition $k_n^{+n}$ ($\mu\text{M}^{-1} \text{s}^{-1}$) | Native monomer removal $k_n^{-n}$ ($\text{s}^{-1}$) |
| --- | --- | --- |
| 1-mer | $(2 \pm 2) \times 10^{-9}$ | N/A |
| 2-mer | $(2 \pm 2) \times 10^{-8}$ | $(2 \pm 2) \times 10^{-7}$ |
| 3-mer | N/A | $(2 \pm 1) \times 10^{-6}$ |

|  |  |
| --- | --- |
| Misfolding rate $k_m$ ($\text{s}^{-1}$) | $(2.1 \pm 0.4) \times 10^{-7}$ |
| Refolding rate $k_r$ ($\text{s}^{-1}$) | $(1.6 \pm 0.4) \times 10^{-5}$ |
| Initial monomer concentration ($\mu\text{M}$) | $9.3 \pm 0.1$ |
| Initial dimer concentration ($\mu\text{M}$) | $1.02 \pm 0.02$ |
| Initial trimer concentration ($\mu\text{M}$) | $0.161 \pm 0.06$ |

**Table S6:**
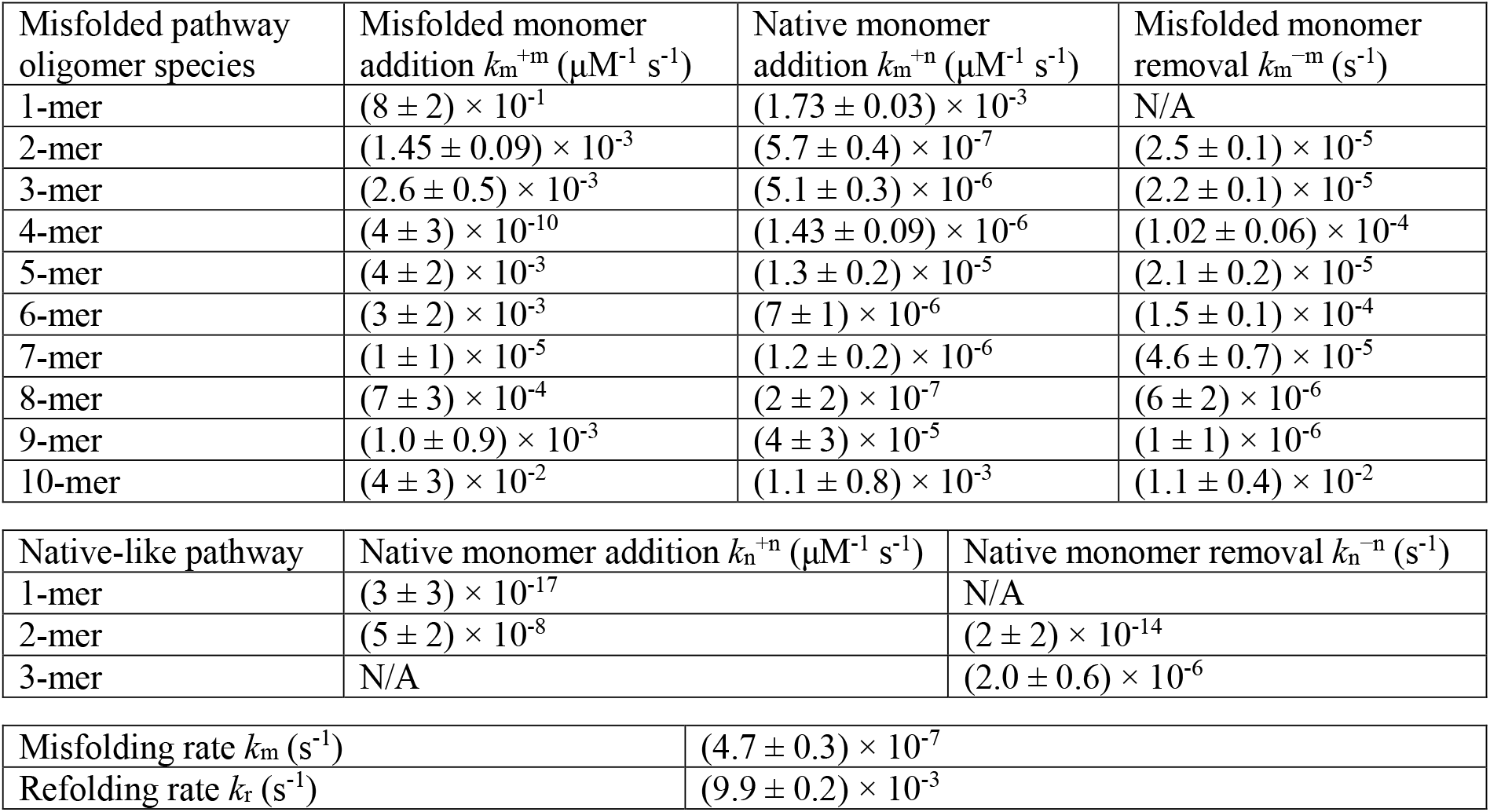
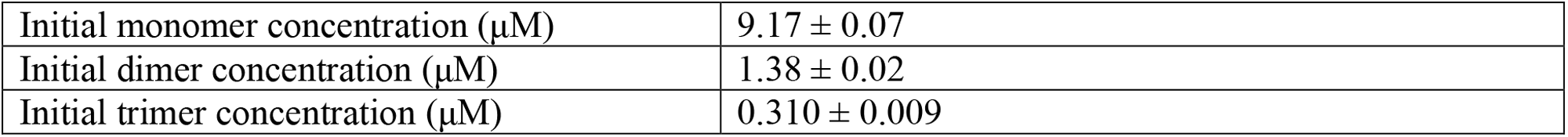
SMMP fit results with methylene blue. Results from fitting SMMP data in Fig. 5B to model 2.

| Misfolded pathway oligomer species | Misfolded monomer addition $k_m^{+m}$ ($\mu\text{M}^{-1} \text{s}^{-1}$) | Native monomer addition $k_m^{+n}$ ($\mu\text{M}^{-1} \text{s}^{-1}$) | Misfolded monomer removal $k_m^{-m}$ ($\text{s}^{-1}$) |
| --- | --- | --- | --- |
| 1-mer | $(8 \pm 2) \times 10^{-1}$ | $(1.73 \pm 0.03) \times 10^{-3}$ | N/A |
| 2-mer | $(1.45 \pm 0.09) \times 10^{-3}$ | $(5.7 \pm 0.4) \times 10^{-7}$ | $(2.5 \pm 0.1) \times 10^{-5}$ |
| 3-mer | $(2.6 \pm 0.5) \times 10^{-3}$ | $(5.1 \pm 0.3) \times 10^{-6}$ | $(2.2 \pm 0.1) \times 10^{-5}$ |
| 4-mer | $(4 \pm 3) \times 10^{-10}$ | $(1.43 \pm 0.09) \times 10^{-6}$ | $(1.02 \pm 0.06) \times 10^{-4}$ |
| 5-mer | $(4 \pm 2) \times 10^{-3}$ | $(1.3 \pm 0.2) \times 10^{-5}$ | $(2.1 \pm 0.2) \times 10^{-5}$ |
| 6-mer | $(3 \pm 2) \times 10^{-3}$ | $(7 \pm 1) \times 10^{-6}$ | $(1.5 \pm 0.1) \times 10^{-4}$ |
| 7-mer | $(1 \pm 1) \times 10^{-5}$ | $(1.2 \pm 0.2) \times 10^{-6}$ | $(4.6 \pm 0.7) \times 10^{-5}$ |
| 8-mer | $(7 \pm 3) \times 10^{-4}$ | $(2 \pm 2) \times 10^{-7}$ | $(6 \pm 2) \times 10^{-6}$ |
| 9-mer | $(1.0 \pm 0.9) \times 10^{-3}$ | $(4 \pm 3) \times 10^{-5}$ | $(1 \pm 1) \times 10^{-6}$ |
| 10-mer | $(4 \pm 3) \times 10^{-2}$ | $(1.1 \pm 0.8) \times 10^{-3}$ | $(1.1 \pm 0.4) \times 10^{-2}$ |

| Native-like pathway | Native monomer addition $k_n^{+n}$ ($\mu\text{M}^{-1} \text{s}^{-1}$) | Native monomer removal $k_n^{-n}$ ($\text{s}^{-1}$) |
| --- | --- | --- |
| 1-mer | $(3 \pm 3) \times 10^{-17}$ | N/A |
| 2-mer | $(5 \pm 2) \times 10^{-8}$ | $(2 \pm 2) \times 10^{-14}$ |
| 3-mer | N/A | $(2.0 \pm 0.6) \times 10^{-6}$ |

|  |  |
| --- | --- |
| Misfolding rate $k_m$ ($\text{s}^{-1}$) | $(4.7 \pm 0.3) \times 10^{-7}$ |
| Refolding rate $k_r$ ($\text{s}^{-1}$) | $(9.9 \pm 0.2) \times 10^{-3}$ |
| Initial monomer concentration ($\mu\text{M}$) | $9.17 \pm 0.07$ |
| Initial dimer concentration ($\mu\text{M}$) | $1.38 \pm 0.02$ |
| Initial trimer concentration ($\mu\text{M}$) | $0.310 \pm 0.009$ |

**Figure S1:**
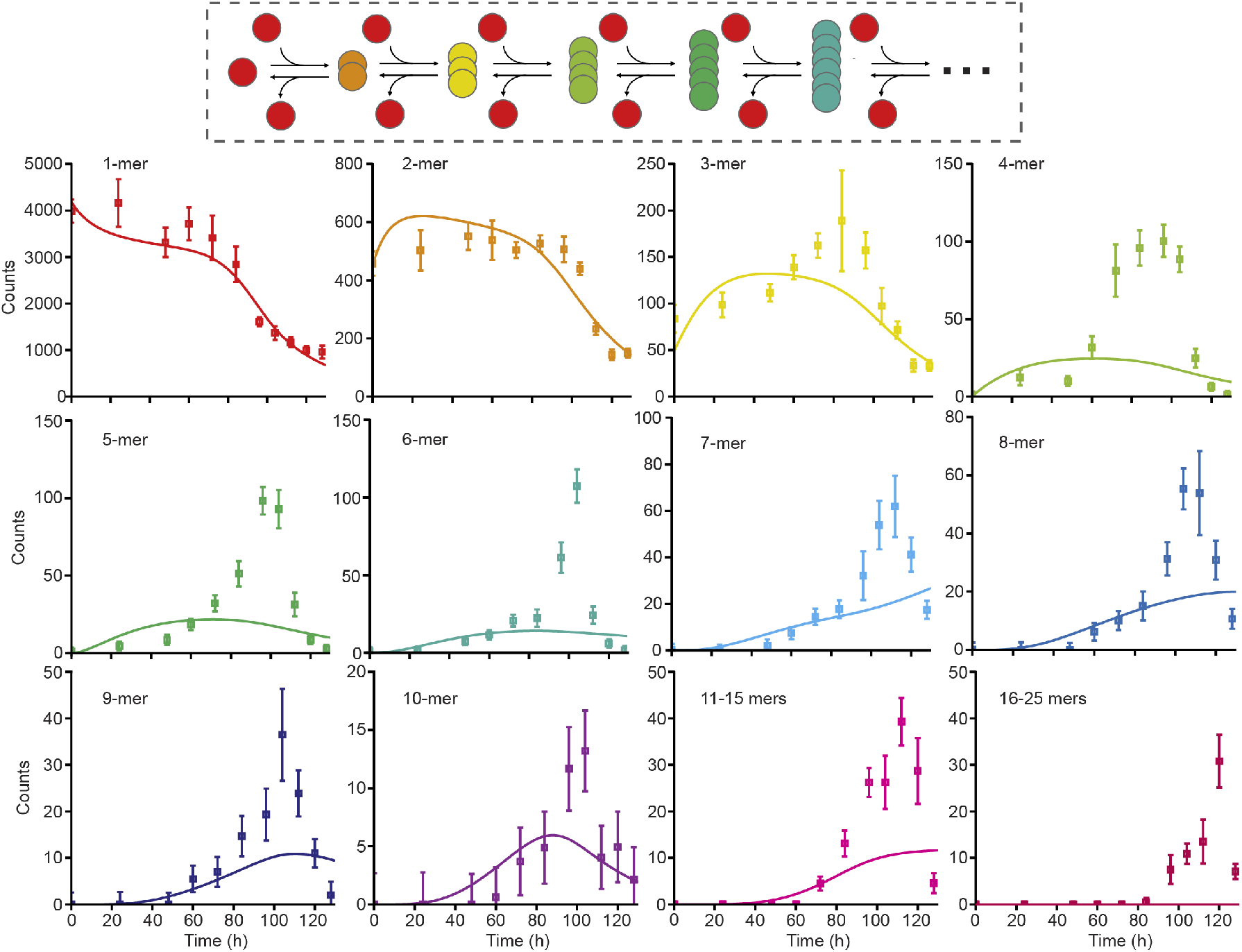
Single-pathway aggregation model does not capture oligomer kinetics. Fitting the aggregation data to a model featuring a single pathway in which all oligomers can grow without limit fails to capture the oligomer population dynamics observed with SMMP.

**Figure S2:**
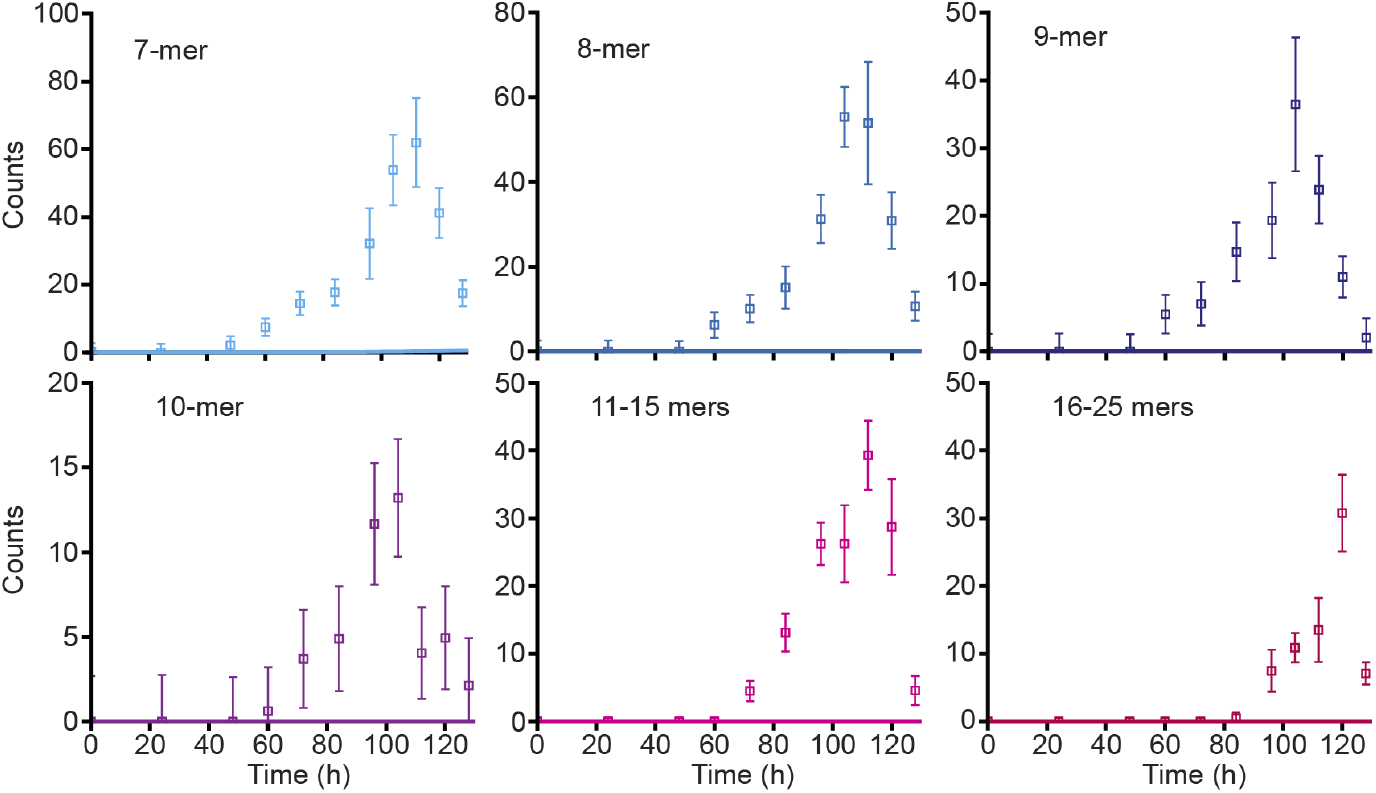
Saturating elongation and fragmentation model from fitting ThT data fails to predict dynamics of larger oligomers observed by SMMP. The model that best fits the concentrationdependent fibrillization data (saturating elongation and fragmentation) predicts very low levels for oligomers from 7-mers to 25-mers, in disagreement with populations observed by SMMP.

**Figure S3:**
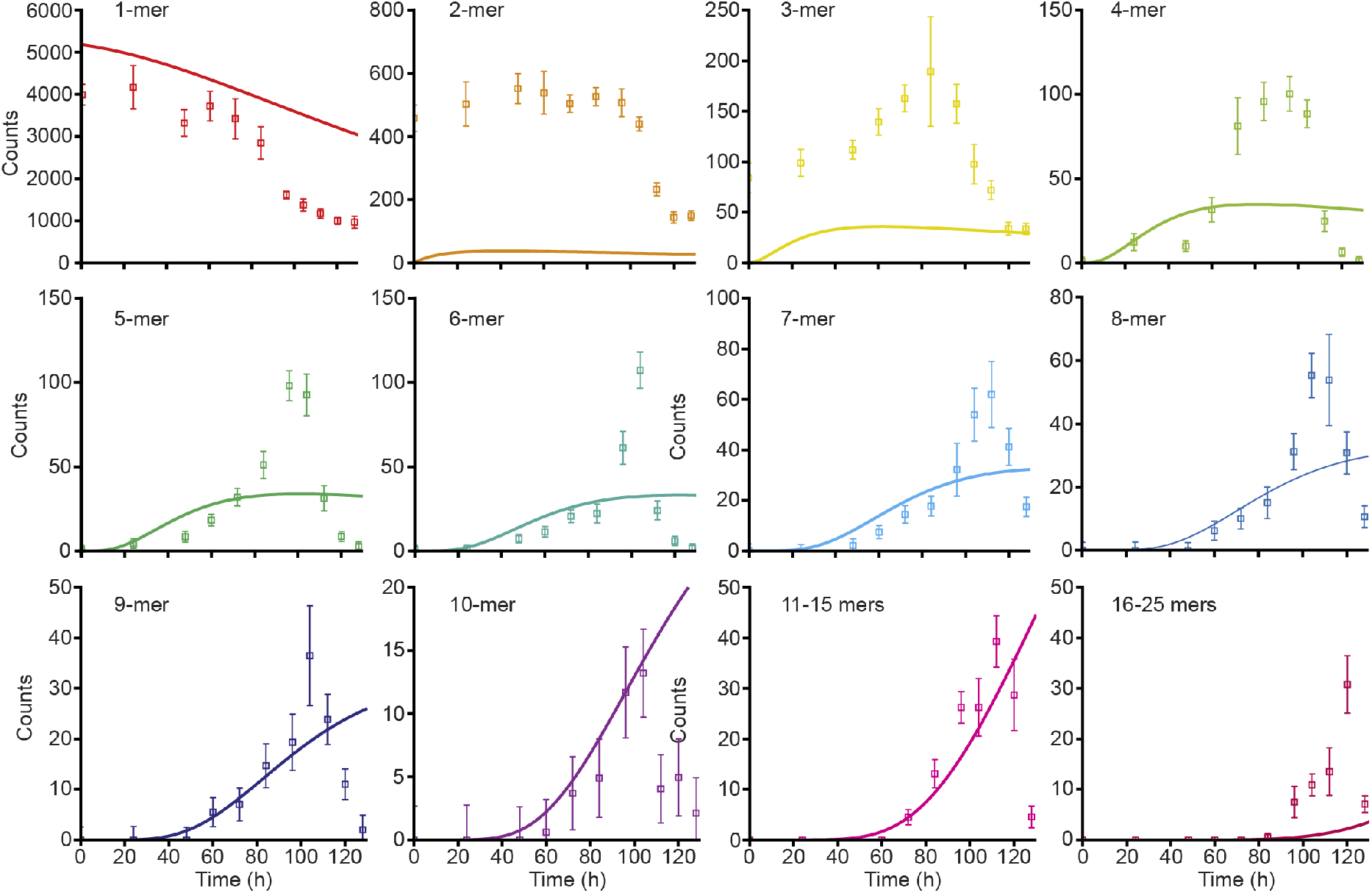
Oligomer populations predicted by fit of ThT data to nucleation-elongation model. Despite its poor fit to the ThT data, the nucleation-elongation model predicts the oligomer populations better than the saturating elongation and fragmentation model. However, the predictions still do not match the populations observed by SMMP.

**Figure S4:**
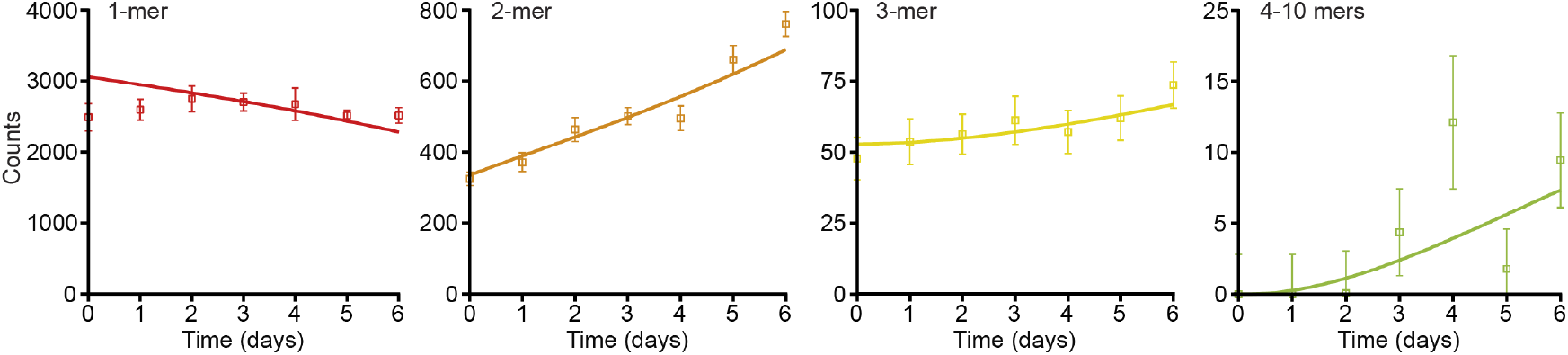
Modeling of oligomer populations in the presence of EGCG. Owing to the overall lack of aggregation, the oligomer populations show only minor changes. The kinetics are fit well by model 2 (solid lines).

**Figure S5:**
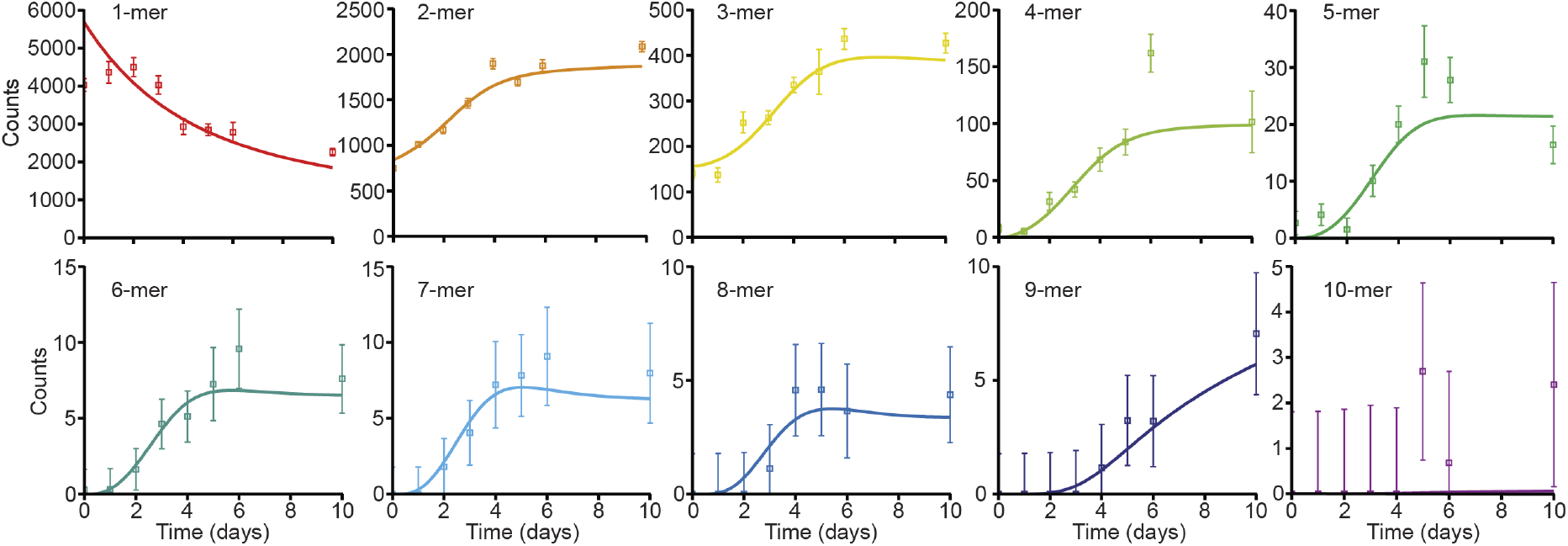
Modeling of oligomer populations in the presence of methylene blue. Small but noticeable amounts of higher-order oligomers are produced in the presence of methylene blue. The kinetics are fit well by model 2 (solid lines).

**Figure S6:**
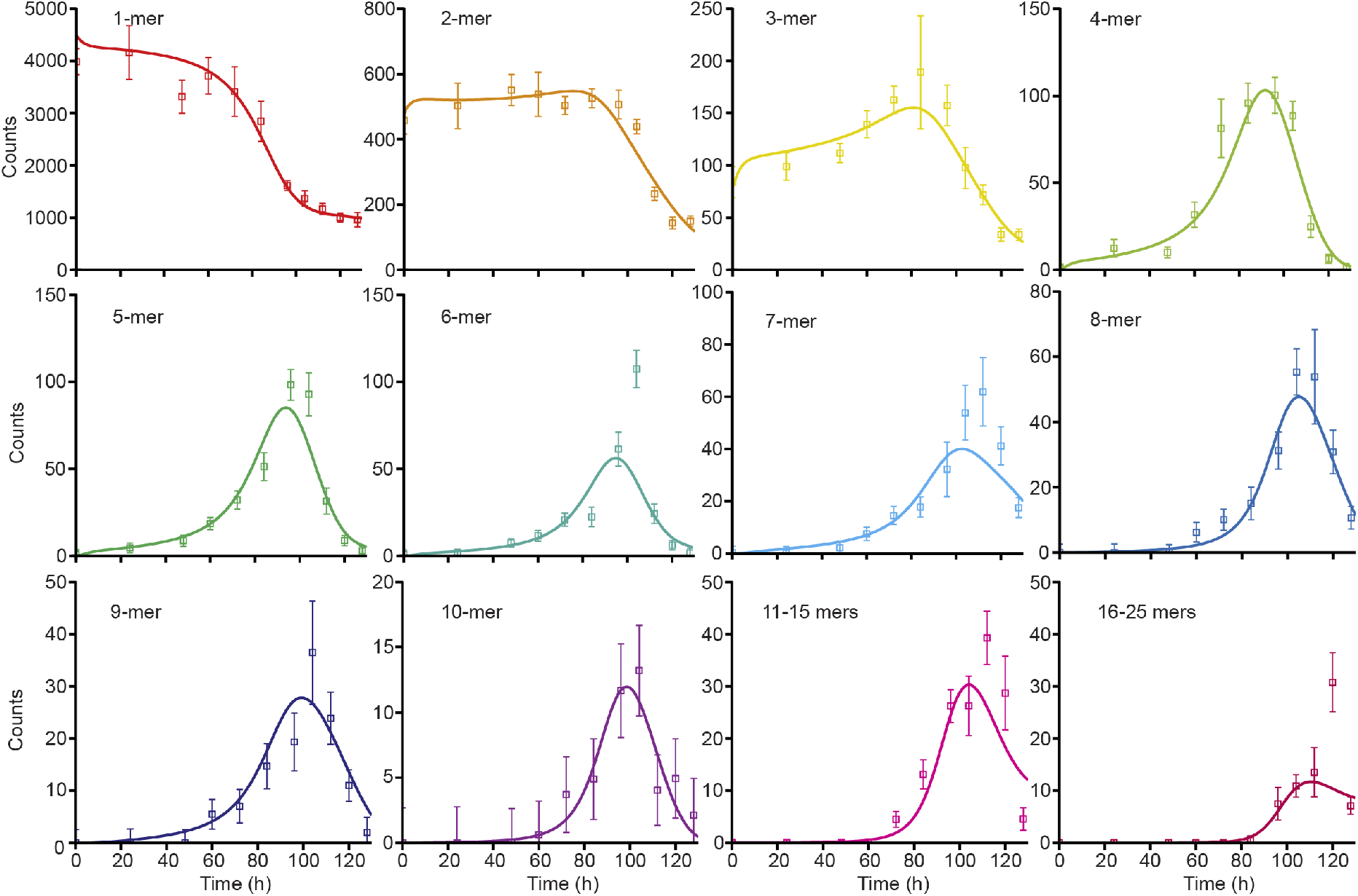
Global fitting of SMMP and ThT data does not affect fit to oligomer kinetics. Jointly fitting both the oligomer populations and the ThT signal in Fig. 3 leads to predictions of oligomer kinetics that are nearly indistinguishable from those obtained by fitting the SMMP data alone.

